# Diverse northern Asian and Jomon-related genetic structure discovered among socially complex Three Kingdoms period Gaya region Koreans

**DOI:** 10.1101/2021.10.23.465563

**Authors:** Pere Gelabert, Asta Blazyte, Yongjoon Chang, Daniel M. Fernandes, Sungwon Jeon, Jin Geun Hong, Jiyeon Yoon, Youngmin Ko, Victoria Oberreiter, Olivia Cheronet, Kadir T. Özdoğan, Susanna Sawyer, Songhyok Yang, Ellen McRae Greytak, Hansol Choi, Jungeun Kim, Jong-Il Kim, Kidong Bae, Jong Bhak, Ron Pinhasi

## Abstract

The genetic history of prehistoric and protohistoric Korean populations is not well understood due to the lack of ancient Korean genomes. Here, we report the first paleogenomic data from Korea; eight shotgun-sequenced genomes (0.7×∼6.1× coverage) from two archeological sites in Gimhae: Yuha-ri shell mound and Daesung-dong tumuli, the most important funerary complex of the Gaya confederacy. All eight individuals are from the Korean Three Kingdoms period (4th-7th century CE), during which there is archaeological evidence of extensive trade connections with both northern (modern-day China) and eastern (modern-day Japan) kingdoms. All genomes are best modeled as an admixture between a northern-Chinese Iron Age genetic source and a Japanese-Jomon-related ancestry. The proportion of Jomon-related ancestry suggests the presence of two genetic groups within the population. The observed substructure indicates diversity among the Gaya population that is not related to either social status or sex.

**Teaser:** 1,700-year-old genomes reveal the genetic diversity of ancient Koreans in the Gimhae region.

## Introduction

Recent studies reveal that East Asian populations have a significant genetic continuity that can be traced back to the Neolithic (*1–8*). Present-day Koreans are an East Asian group with distinct genetic and linguistic traits. The Korean language is often classified as an isolate, with no other closely related modern languages (*9*). A recent study of 1,094 present-day Korean genomes indicates that they are a genetically homogenous group (*10*). This apparent homogeneity may be the outcome of isolation during the last centuries due to genetic drift. Additionally, Korean maternal and paternal markers hint towards multiple lineages forming the present-day Korean population through admixture events in the past (*10, 11*). Modern Korean genomes can be modeled with Bronze Age populations of the Liao River (China), as well as ancestry associated with ancient individuals from the Japanese archipelago, and Devils’ Gate genomes in Northern Asia (*1*).

Agricultural practices and complex societies emerged in Korea during the Bronze Age (1400-300 BCE) (*12*). Both the genetic study of rice (*12*) and archeological research(*13*) suggest that rice, and crops including millet and various legumes, were introduced and subsequently intensively used in the Korean peninsula around 850-550 BCE (*13*). Up to this day, the exact location from where the rice was introduced in the Korean peninsula is controversial, however, there is a consensus that rice cultivation spread from the Korean peninsula to the Japanese archipelago (*14*). After agriculture, iron production arrived from China to Korea around the 4th century BCE (*15, 16*), and further spread from Korea to Japan during the Yayoi period (10th century BCE-3rd century CE) (*17*). However, it is unclear whether the spread of iron technology in the Korean peninsula was accompanied by human migration or transferred only via trade and cultural diffusion. The Three Kingdoms (TK) period of Korea (4th-7th century CE) was an era of extensive development of iron technology and trading with neighboring populations. During the TK period, the Korean Peninsula was ruled by several states (*15*) including Goguryeo, Baekje, and Silla. The Gaya confederacy was the last independent territory that lasted until the 6th century CE before being assimilated by the neighboring Silla (*18*). Although absorbed by Silla eventually, Gaya had the most developed iron production and trading infrastructure during the early TK period, including exchange of goods, and possibly also the movement of people, with the Wa (Wae) inhabitants of the Japanese archipelago and northern China (*19*). This suggests that Gimhae, the political center of Gaya, was an important trading centre and as such its populace may have included a level of cosmopolitanism as has been reported, for example, in a recent genetic study of the ancient inhabitants of Rome (*20*).

At the end of the 3rd century CE archaeological sites from the Gimhae area underwent changes in funerary rituals that are marked by wooden-chambered burials entirely replacing and the preceding burials characterized by the stone-lined pit tombs, and jar coffin tombs (*21*). Another element that was introduced in this period was a mortuary practice that involved human sacrifices and the inclusion of various grave goods such as weapons, mirrors and other items made of bronze and iron as well as severed heads of livestock, that accompanied the deceased (the main burial) (*19*). The most important excavated center of the Gaya culture is a massive burial complex of rulers in Daeseong-dong in Gimhae, dated 1st-5th century CE. It covers an area of 3,700 m^2^ and consists of 219 tombs, 69 of which are complexes with multiple burials including human sacrifices and various objects such as pottery, iron armors and items related to archery. More modest burials existed near shell mounds in the same Gyungsang province, in areas such as Busan and Gimhae throughout the TK period. Most of these mounds did not include human sacrifices, but often included a family grave or a single burial (*19*) such as a female child skeleton from the Yuha-ri shell mound (AKG_3420).

The TK period was a major formative era in the protohistory of Korea and most likely had a major impact on the genetic structure of modern Korean populations. However, to date there are no paleogenomic data for any prehistoric or protohistoric Korean populations. Here, for the first time, we assess the genetic structure of eight TK period individuals from Gaya confederacy in relation to their social status (inferred from mortuary contexts) and the relationship between the TK individuals and present-day Koreans.

## Results

We screened 27 petrous bones and teeth obtained from 22 individuals from two archeological sites in Gimhae city, Gyeongsangnam-do: the Daeseong-dong and Yuha-ri, both dated to 4th–5th century CE. The libraries of eight individuals had over 7% of sequencing reads aligning to the human reference genome (range from 7.3% to 68.9% (Table S1)). The sequencing yielded coverage depths between 0.7× and 6.1× (Table 1). All seven Daeseong-dong individuals were associated with specific mortuary practices which are linked to social status, specifically, main burials, and human sacrifices. A single grave of a child (AKG_3420) from the Yuha-ri shell mound (Table 1, Fig. 1B, Table S1, Fig. S1-S3) could not be associated with social status due to the lack of mortuary elements. All eight genomes exhibited typical deamination patterns (*22*) at read ends (Table 1, Table S2, Fig. S4). The deamination rates in the last bases of 5’ and 3’ ends ranged between 22% and 33%. Five of the individuals were identified as female and three as male based on sex chromosomes and autosomes depth coverage ratios. There was no clear identifiable familial relationship among the sequenced individuals based on the likelihood method lcMLkin (*23*). The lack of kinship evidence was supported by no signs of inbreeding observed from Runs of Homozygosity (RoH) analysis using a software based on pseudo haploid data (*24*) (Fig. S5).

**Fig. 1:**
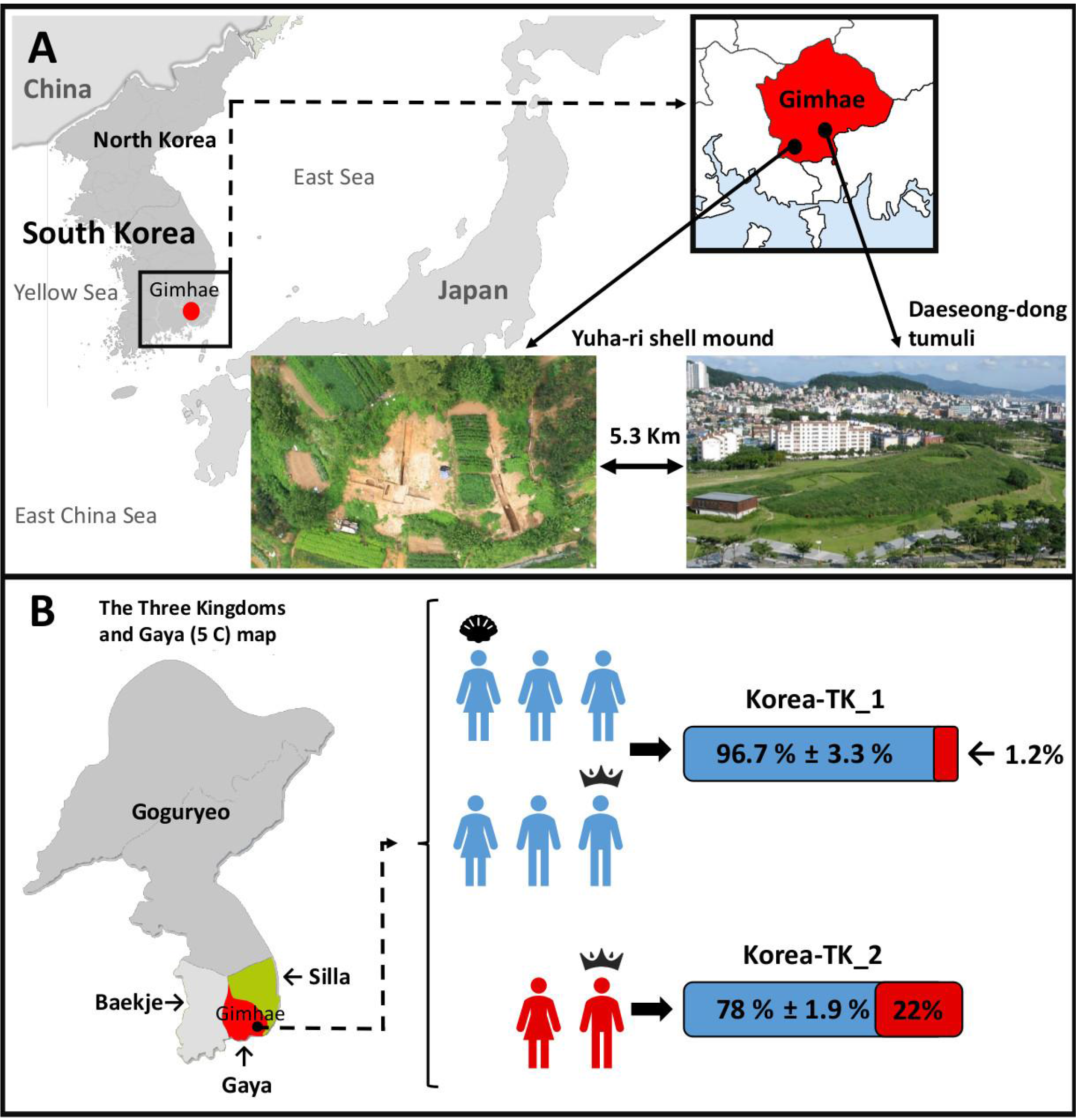
Three Kingdoms Period genomes from Korea. **A)** Site description: geographical location and images of the sites of Daeseong-dong and Yuha-ri. The image of the Yuha-ri shell mound aerial view, and the panoramic view of Daeseong-dong tumuli show a general perspective of the archeological sites. **B)** Genomic composition of the AKG individuals: We present the eight individuals classified into the two identified genetic groups based on qpAdm analyses: TK_1 (blue icon), and TK_2 (red icon), the percentage bar graph shows the genetic ancestry proportions of: North-China Iron Age ancestry (blue), Jomon-related ancestry (red). Individuals in panel B are also classified by the archeological context. Crown depicts a main burial, and a shell depicts the shell mound burial, the samples without a distinct marking are human sacrifices.

**Table 1:** Main descriptive information of the analyzed individuals.

| Individual | Cluster | Date | Typology | Coverage |
| --- | --- | --- | --- | --- |
| AKG_10203(♂) | Korea-TK_2 | 400-450 CE | Daeseong-dong-Burial | 1.58× |
| AKG_10204(♂) | Korea-TK_1 | 400-450 CE | Daeseong-dong-Sacrifice | 0.79× |
| AKG_10207(♀) | Korea-TK_2 | 375-400 CE | Daeseong-dong-Sacrifice | 6.11× |
| AKG_10209(♀) | Korea-TK_1 | 425-450 CE | Daeseong-dong-Sacrifice | 1.78× |
| AKG_10210(♀) | Korea-TK_1 | 425-450 CE | Daeseong-dong-Sacrifice | 2.72× |
| AKG_10218(♂) | Korea-TK_1 | 400-470 CE | Daeseong-dong-Burial | 0.64× |
| AKG_3420(♀) | Korea-TK_1 | 400 CE | Yuha-ri shell mound | 3.20× |
| AKG_3421(♀) | Korea-TK_1 | 300-350 CE | Daeseong-dong-Sacrifice | 0.74× |

We generated two base-calling sets with these genomes. First, we called pseudo haploid positions of the 1,240k dataset (*25*), a standard call-set that is used in most of the downstream analyses involving ancient samples. Second, we generated imputed diploid calls that were used to identify haplotype-based connections with present-day populations, following previously described methodologies (*26*).

### Uniparental markers

All the samples have typical East Asian mitochondrial DNA (mtDNA) haplogroups (Table S3) : D (n=5), B, F, and M when we determined the haplogroups from the consensus sequences obtained from Schmutzi (*27*) with HaploGrep2 (*28*). All the identified mtDNA haplogroups prevail among present-day Koreans (*9, 11*). The most common Korean mtDNA haplogroup is D4, which we identified in four of the individuals. This haplogroup is also common among Japanese Yayoi farmers, while absent in Jomon (*29*). Out of the three male individuals in this study, we successfully called Y chromosome haplogroups for two: AKG_10203 (D1a2a1) and AKG_10204 (O1b2a1a2a1b1). The third male, AKG_10218, ecould not offer fine resolution, but was nonetheless able to be assigned an O haplogroup. Haplogroup O is the most common Y haplogroup shared by more than 73% of Korean males (*9*), and haplogroup D is more common in the present-day Japanese population (*30*).

### Population genetics based on Principal Component Analysis, qpWave, and ADMIXTURE

We focused on the Human Origins (HO) (*31*) panel of SNPs and compared those for the eight genomes with the rest of East Asian modern and ancient diversity. A Principal Component Analysis (PCA) performed with smartPCA showed the presence of three differentiated genetic groups in ancient East-Asia: 1) the Amur River populations, 2) northern China, and 3) southern China and Taiwanese populations (Fig. 2A, Fig. S6, Table S4-S5). The eight individuals from the TK fall clearly within the diversity of the East Asian samples, especially present-day Koreans and Japanese (Fig. 2A). However, two samples (AKG_10203 and AKG_10207), clustered rather closely with present-day Japanese and closer to Japanese Jomon individuals (Fig. 2A), a pattern which distinguishes them from the other six individuals which cluster with present-day Koreans.

**Fig. 2:**
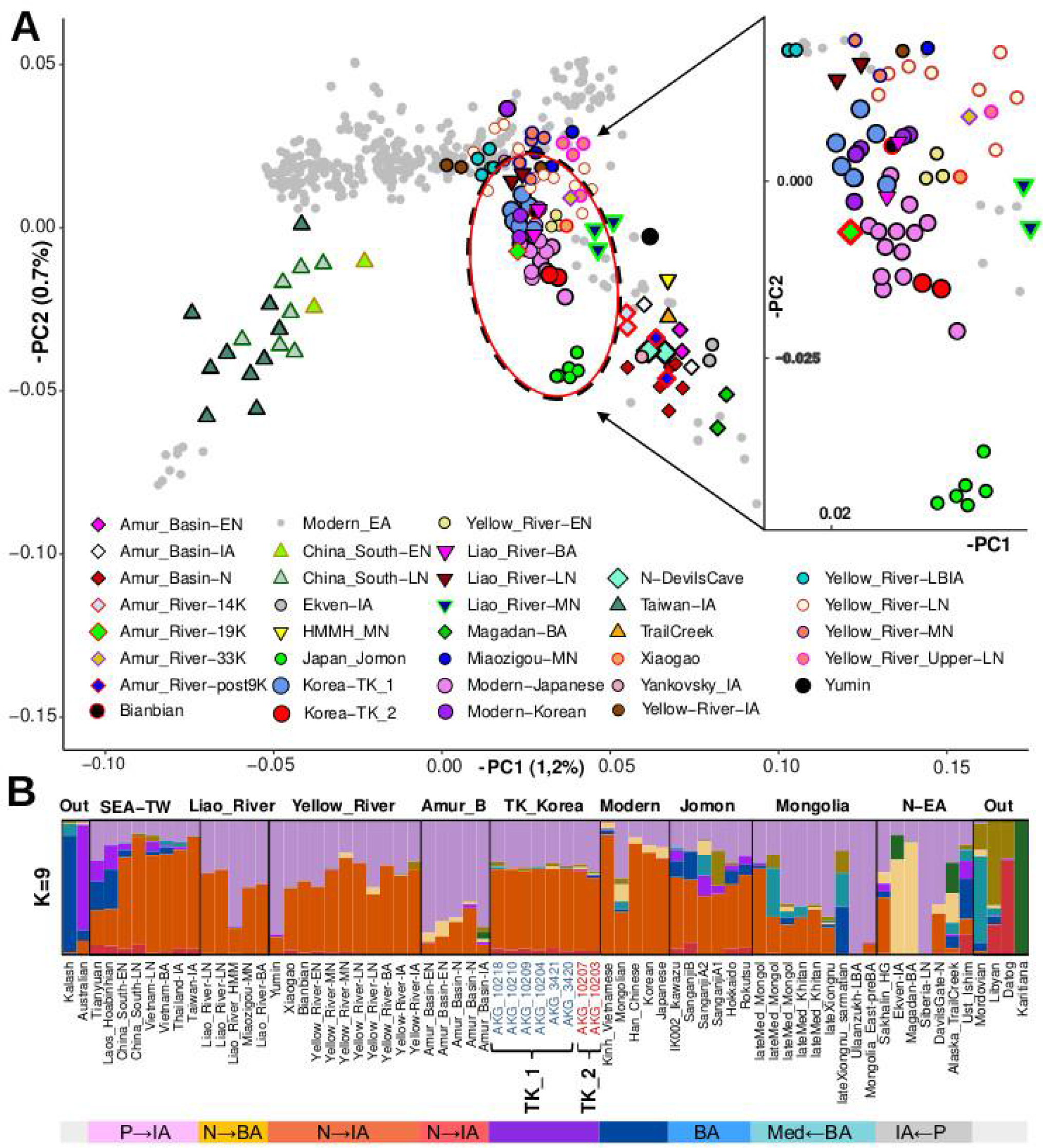
TK period Korean genomic context: A) Principal components analysis performed with present-day East-Asian populations and 96 projected ancient individuals, including eight TK period Korean samples. B) ADMIXTURE plot (K=9) of the most representative populations used in the analyses show a shared genomic profile across ancient and modern East Asians, including TK Koreans and present-day Koreans. Outgroups (2 on the left and 4 on the right) show populations where minor components from East Asia appear as major. The letters and arrows in the colored blocks indicate the chronological order of the samples presented: P->IA Paleolithic to Iron Age; N->BA Neolithic to Bronze Age; N->IA Neolithic to Iron Age; BA - all samples belong to Bronze Age; Med<-BA Medieval to Bronze Age, etc.

Next, we further explored the ancestral compositions of the eight individuals using a pairwise qpWave analysis. The main aim is to identify individuals that might not form a clade with the others, based on a specified set of outgroup populations. When using a set of ten outgroup populations, which includes ancient Japanese Jomon populations, most individuals formed a clade with each other. However, in contrast, the two aforementioned individuals, AKG_10203 and AKG_10207, formed a clade only with each other, which is related to their shared Jomon-related ancestry. We then separated our eight TK individuals into two groups: (a) the two outliers (Korea-TK_2), and (b) the remaining six individuals (Korea-TK_1). This grouping is also supported by the diploid call analyses, which shows that one individual (AKG_3421) has an intermediate position between both clusters and had a *p*-value between 0.01-0.05 in the qpWave clustering analysis (Fig. S7). Despite these facts, the *f_4_* statistics *f_4_*(Mbuti, Japan_Jomon; AKG_3421, Korea-TK_1_individual) of each individual within Korea-TK_1 has a z-score of Z<|1.75|, which supports the inclusion of AKG_3421 within Korea-TK_1. The clustering in the PCA plot could not be linked to the mortuary status as both Korea-TK_1 and Korea_TK_2 have groups consisting of both main burial and sacrifice burials (Table 1).

We further explored the genetic structure of the Gimhae Korean individuals with ADMIXTURE analysis, which included 2,283 present-day and ancient individuals (Fig. 2B, Fig. S8) and a pruned SNP set of the Human Origin (HO) dataset (extended methods). At K=9, the Korea-TK individuals could be represented by two major genetic components shared with the Neolithic Devils_Gate (Russian Far East) genomes (Fig. 2B). One is mainly characteristic of Siberian Late Neolithic genomes (light purple), and the other is a major genetic component that is commonly present in most ancient and present-day East Asians, and is especially characteristic of ancient southern Chinese, southeast Asians, and modern Kinh Vietnamese (orange). Overall, the Korea-TK_1 and Korea-TK_2 individuals have a genomic composition that closely resembles Yellow River Bronze and Iron Age individuals and Bronze Age Liao River populations. The minor components found in TK period Koreans, especially in Korea-TK_2, are shared with Jomon. AKG_10203 and AKG_10207, the Korea-TK_2 samples, clustered together in the PCA and qpWave analyses. In the ADMIXTURE modelling, they have a genomic profile that differs slightly from each other in its minor components. However, in both cases the minor components can be linked to the components abundant in Jomon and those that are found only as minor components among southern prehistoric populations from southern China, Thailand, Vietnam, and adjacent regions. Notably, the Jomon-related ancestry component is absent among present-day Koreans, suggesting that the population at Gaya was more diverse than nowadays Korean population. Additionally we suggest that Korea-TK_2 may not have contributed substantially to the genetic structure of present-day Koreans.

### The origin of the Korean Three Kingdoms Period population using f-statistics

We computed outgroup *f_3_* statistics, which measures the amount of shared genetic similarity between populations. We tested both groups, Korea-TK_1 and Korea-TK_2, and compared them to ancient and present-day populations (Fig. 3A-3D, Fig. S9-10, Tables S6-S7). Korea-TK_1 showed major affinities to ancient eastern Chinese coastline populations, while Korea-TK_2 appeared to be the closest to Jomon. A heatmap based on *f_3_* comparisons (Fig. S11) confirmed that the TK period Korean population shared affinities with eastern Chinese populations. When compared to various present-day Asian populations, present-day Koreans and Han Chinese were the closest groups to Korea-TK_1 while the Japanese were the closest group to Korea-TK_2 (Fig. 3A-3B, Table S6).

**Fig. 3:**
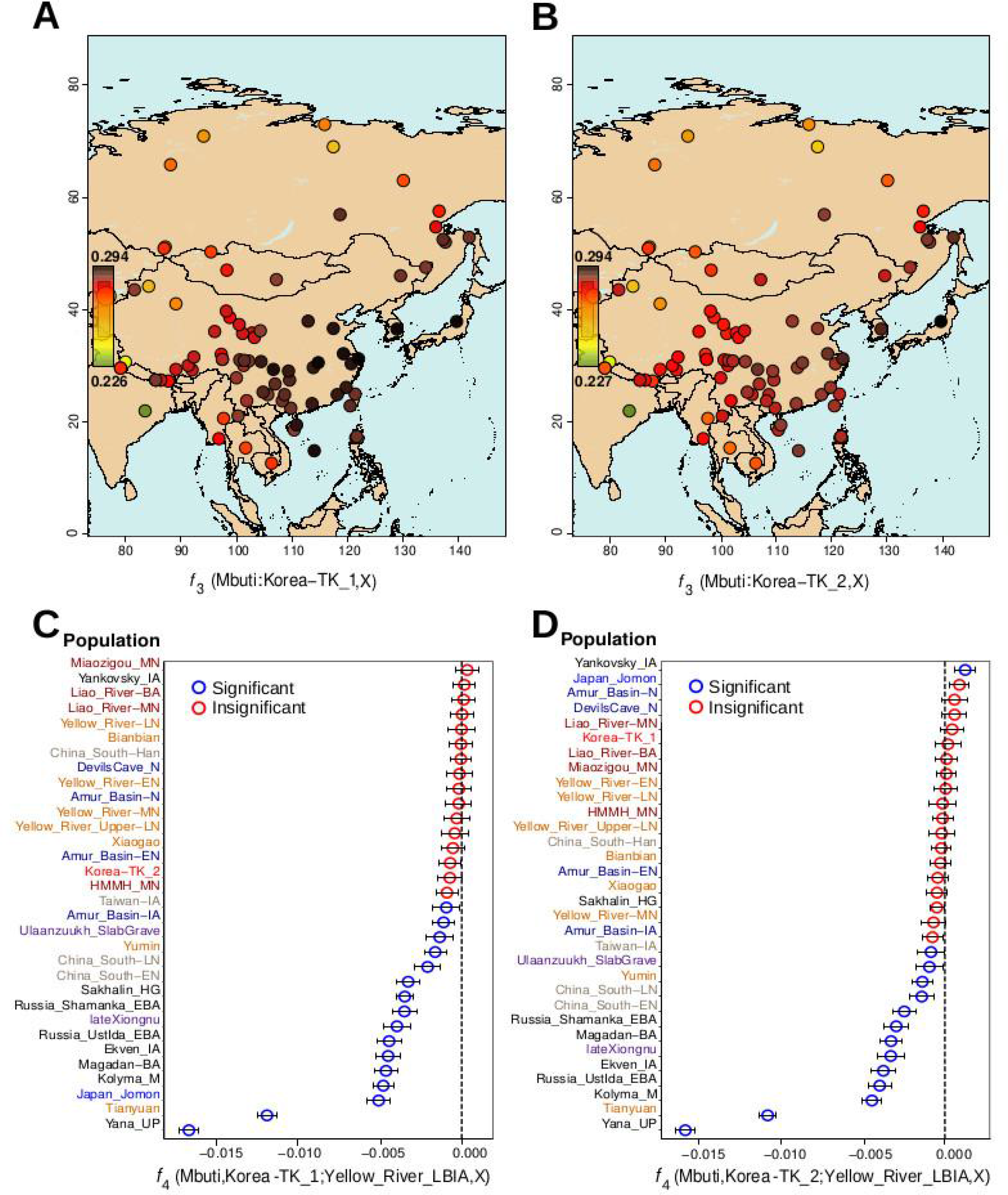
*f*-statistics: A-B) *f*_3_-statistics of present-day Asian populations with ancient Koreans in the combination *f*_3_(Mbuti; Korea-IA,X). The figure shows the connection between East Asian populations and Korea-IA populations. The shared genetic drift between Korea-TK_2 and Japan_Jomon is higher than the Korea-TK_2 and Japan_Jomon. C) qpWave plot: the qpWave analysis shows the presence of two differentiated clusters. *f_4_*-statistics: Korea-ITK_1 is associated with ancient populations from Eastern China, while Korea-TK_2 has the closest affinity with Amur Basin and Japan_Jomon populations. Colors in the sample name denote the region as follows: Dark red – Liao River, Orange – Yellow River, Ivory – Southern China and Taiwan, Dark blue – Amur Basin, purple – Mongolia, black – Russia (Russian Far East, Siberia). Additionally, Red denoted Korea-TK samples, Blue denotes Jomon.

**Fig. 4:**
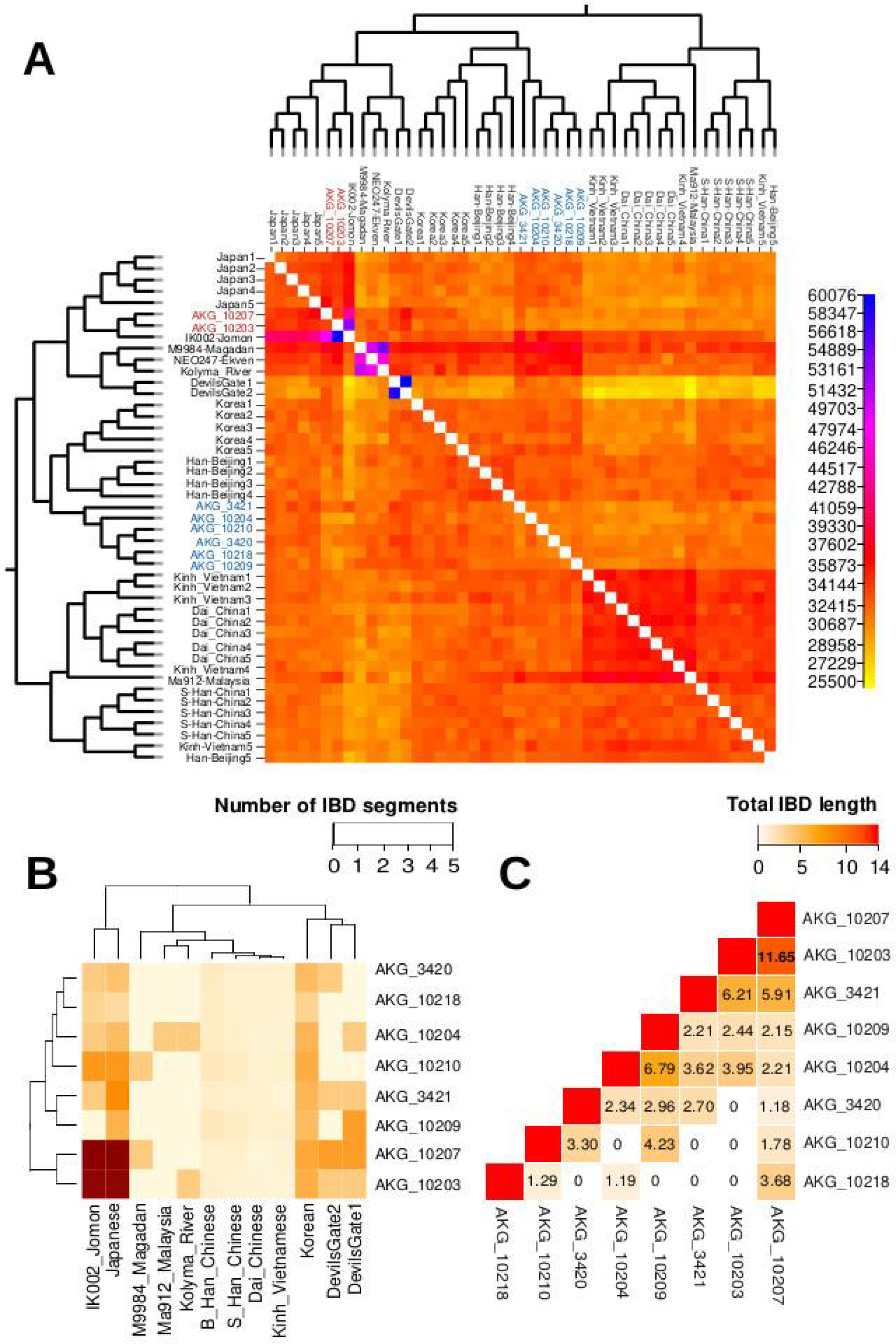
Diploid calls analyses. A) fineSTRUCTURE. The plot shows that Korea-TK_2 clusters with Japanese and IK002_Jomon individuals, while Korea-TK_1 shows connections with Han Chinese and Korean individuals. B) Shared IBD segments (>1 cM). Korea-TK_2 shows the highest number of shared IBD with Japanese, while Korea-TK_1 shows connections with both Koreans and Japanese. C) IBD shared among the pairs of TK individuals; even though none of the pairs indicate familial relationship, it shows the highest IBD sharing between the TK_2 individuals, namely AKG_10203 and AKG_10207.

Based on *f_4_*(Mbuti, Korea-TK; Pop A, Pop B) statistics comparisons (Fig. 3C-3D, Table S8, Fig. S12), we observed once again that Korea-TK_1 individuals were related to Bronze Age populations from the Liao and Yellow rivers in Northern China. The comparative statistic *f_4_*(Mbuti, Korea-TK_1; Yellow-River_LBIA, Liao-River_BA) yielded statistically nonsignificant results (Z<3). Korea-TK_2 individuals, however, had the highest values with Yankovsky_IA and the Japanese Jomon (0.004271 and 0.003897, respectively, in combination with Yellow_River_LBIA).

### Lineage analyses based on ancestry composition

We applied qpAdm in order to investigate the overall ancestry composition of the two groups and to confirm the observed Jomon-related ancestry, to assess the best fitting model. Applying a 2-step model competition approach, Korea-TK_1 could only be modelled as composed by an average of 97±1% of ancestry related to Yellow_River_LBIA and an average of 3.0±1% related to Japan_Jomon (P=0.472) or Japan_Jomon IK002 (p=0.495), whereas Korea-TK_2 required an average of 20.5±1.75% of Jomon-related ancestry (Fig. 1B, Table S9). These results suggest that Korea-TK_2 had ∼20% Jomon-related ancestry, and also points to the presence of Jomon-related ancestry in Korea-TK_1, albeit in much smaller proportions (about 1%). When we performed the same qpAdm modeling on an individual basis, we found that four of the six individuals in the Korea-TK_1 cluster could be modelled with a single source, typically Yellow_River-BA or Liao_River-BA, whereas individual AKG_10218, required between 4.0±1.8% and 7.3±4.4% ancestry related to either the Japanese Jomon or Ekven_IA, respectively, as a second source of ancestry (Table S10, Fig. S13-S14). In turn, individual AKG_3421 required 11.0±2.9% Jomon-related ancestry (p=0.765) (Table S10, Fig. S13-S14), although this model might be overrepresenting Jomon-related ancestry in this individual, ( Z<|1.75|) which indicates nonsignificant affinity to Japan_Jomon when compared to the other Korea-TK_1 individuals.

We attempted to estimate the dates of admixture events identified through the qpAdm analyses using DATES (*32*). We derive an estimate that the Korea-TK_2 source population carrying the Jomon genetic component, was admixed about eight (+/- 3.8) generations prior, before the Korea-TK_2 individuals lived (*32*) (Table S11, Fig. S15). We ran Treemix to graphically represent the relationship with ancient and present-day populations, (Fig. 2B, Fig. S16-S17) with our TK period samples. The best fitting model for Treemix was using seven migration events which showed that Korea-TK_2 appeared admixed with the Jomon population while in Korea-TK_1 Treemix did not pick up a signal of significant shared ancestry (Fig. 5).

**Fig. 5:**
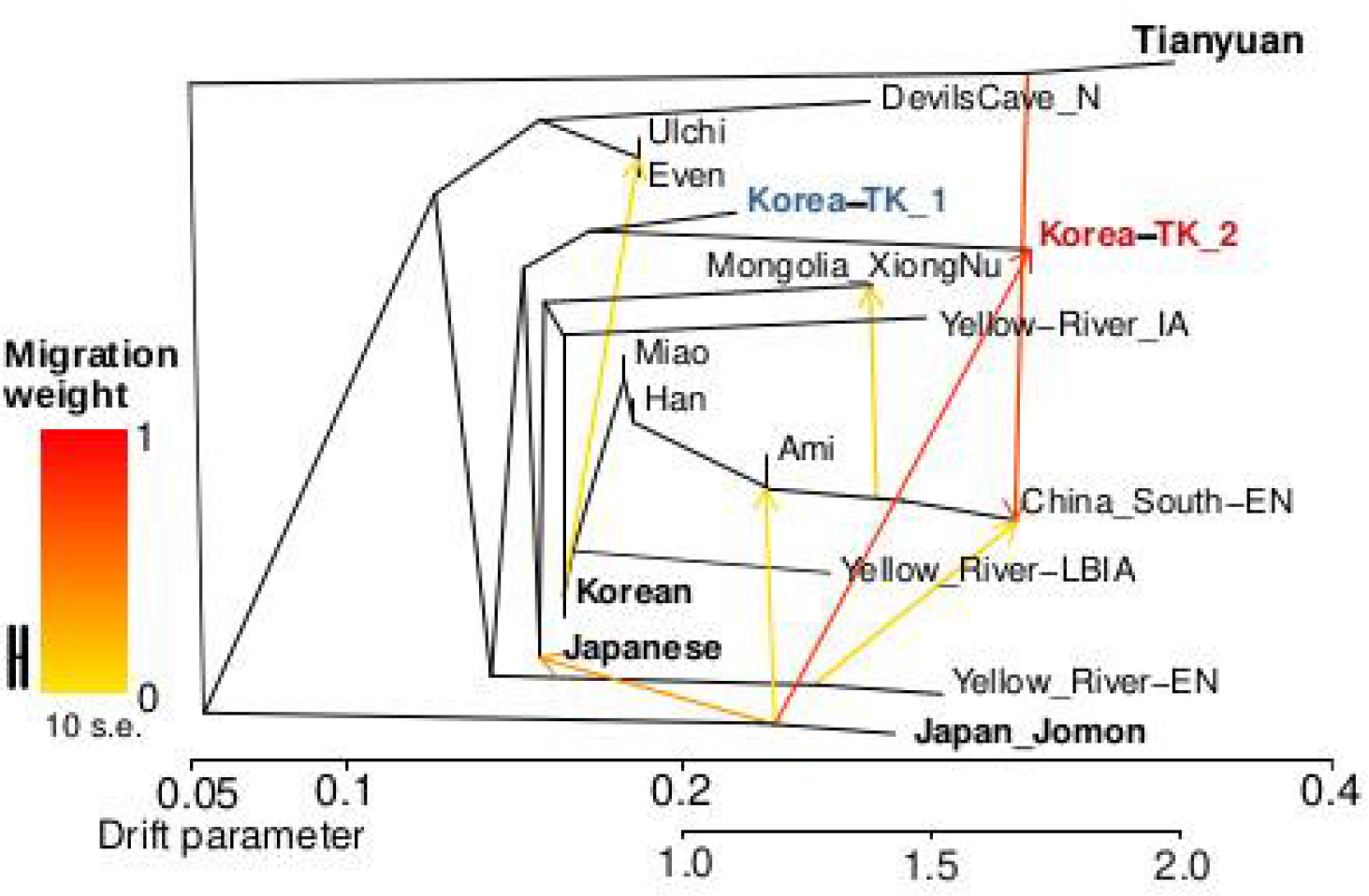
Treemix plot using seven migrations. Korea-TK_1 and Korea-TK_2 groups share the same genetic origin and close relations to present-day Japanese and Koreans, while Korea-TK2 additionally shows genetic ties with ancient Jomon.

Next, we applied qpAdm to model the present-day Koreans using the same ancestry sources that were used for modelling ancient Koreans, including the two ancient Korean groups as separate possible sources. We found that the simplest working models were with a source related to Yellow_River-LBIA (p=0.072) (Table S12). However, when present-day Koreans were modelled using each TK period Korean individual as a source, at least three out of the six individuals from Korea-TK_1 were sufficient enough as a single source (0.099<p<0.908) (Table S12).

Interestingly, while the admixture pattern for the genome of individual AKG_10218, a male, was not different from the models for the Korea-TK_1 individuals, the pattern for his X chromosome SNPs was one that matched a northern Chinese maternal ancestry and lacks the Siberian ancestry associated with Devils_Gate genomes (Fig. S18). This result suggests a differential origin of both parents which is the outcome of an admixture in the region during the TK period.

### Genetic and phenotypic continuity

The results obtained with fineSTRUCTURE and identity-by-descent (IBD) analyses, which were based on the imputed genotypes (see Methods), both show a clear separation between Korea-TK_1 and Korea-TK_2 groups. FineSTRUCTURE suggested Korea-TK_1 individuals were equally related to present-day Koreans and Japanese, a result which was also supported by the higher number and length of IBD blocks that each of the Korea-TK_1 individuals shared with these populations. In regards to Korea-TK_2, the individuals have close genetic affinity to present-day Japanese and the Japanese Jomon from Ikawasu IK002 (Fig. 4A, 4B, Table S13-S14). When compared to the other Korea-TK_1 individuals, the Korea-TK_2 individuals also have a longer length and a higher frequency of shared IBD blocks with present-day Japanese (Fig. 4C). Using a commercial individual-based ancestry prediction service (https://parabon-nanolabs.com), we confirmed that the genetic profile of Korea-TK_1 provides the best match for present-day Korean ancestry while for Korea-TK_2 individuals the closest match was the Japanese Ryukyuan people. Interestingly, the pairwise IBD sharing based on pseudo haploid calls confirmed the absence of close familial relationship among the individuals. However, the two TK2 individuals appeared to share significantly (at least twice) more IBD with each other compared to any other pairing (Fig. 5C).

As significant similarity in SNP profiles and IBD blocks indicate phenotype sharing, we selected 160 phenotypic variants to estimate whether Koreans have changed phenotypically in the last 1,700 years. The small sample size did not enable us to draw any statistically significant inferences, and thus we limited our analysis to presence/absence of the relevant variants. Based on our eight samples, we conclude that all of the relevant variants present in the modern Korean population were already established in the TK period at least 1,700 years ago, showing a deep genetic continuity (Table S15). Some East Asian traits were detected among all the analyzed individuals including dry earwax, no body odor (SNP rs17822931) (*33*), lactose intolerance (rs4988235) (*34*), and no homozygous alleles associated with excess sweating (*35*). All these traits are also common among present-day Koreans. Seven out of eight individuals carried canonical Asian hair straightness/thickness associated with the *EDAR* variant (rs3827760) (*36*) (except AKG_10207); which is homozygous in six of them (Table S15). The HirisPlex system (*37*), a phenotype prediction protocol, predicted all the individuals had brown eyes and most likely black hair and variation in skin tones ranging from intermediate to darker (Tables S16-18), a result which is also in agreement with the phenotypes of present-day Koreans.

Surprisingly, the genetic makeup of the eight Gaya Korean samples also reflected common present-day Korean healthcare and aesthetic concerns: all individuals have multiple (four or more) ‘susceptibility-to-myopia-risk’ alleles. Myopia is highly prevalent in Korea, reaching values of 95% in adult males from Seoul (*38*). In several individuals, we detected variants associated with alcohol flush reaction (*39, 40*). Also, all the samples had on average at least one homozygous balding risk allele or nine androgenic alopecia risk variants (*41*). Sample AKG_10204 has five of these homozygous risk alleles. The eight genomes also capture the diversity of SNP profiles related to eyelid morphology, hair curliness, and other phenotypic features (Fig. 6, Fig. S19). The female sample AKG_10207 from Korea-TK_2 group stood out with a more Jomon-like phenotypic SNP profile, mainly due to homozygosity of reference allele “A” in the *EDAR* gene (rs3827760) (Table S18). Even though it is hard to assess hair morphology, in addition to reference allele homozygosity in *EDAR*, AKG_10207 had multiple SNPs related to hair curliness, three of them homozygous, suggesting she might have had more than slightly wavy hair, while the other samples suggested most likely had straight hair.

**Fig. 6:**
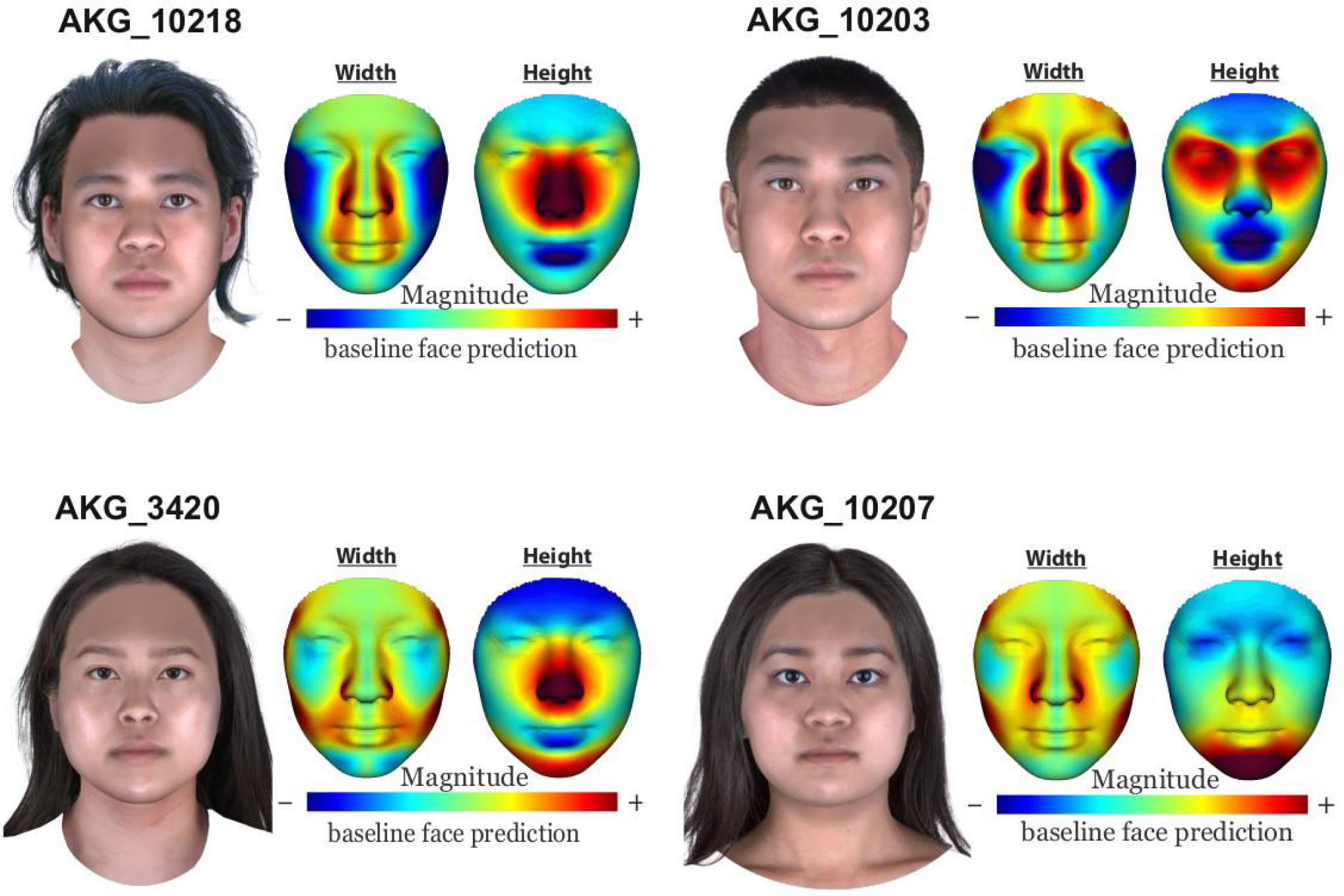
Facial predictions of TK period individuals. Face reconstruction of the Korean TK period individuals AKG_10218, AKG_10203, AKG_3420, AKG_10207 shows a diversity in phenotypic features commonly observed in present-day Koreans. Face morphology differences are emphasized relative to a baseline face prediction made using sex and ancestry. The facial reconstruction has been performed with the imputed calls dataset.

All the phenotypic features assessed by HirisPlex phenotype prediction protocol and our selected forensically relevant facial morphology-related SNPs were independently validated by the Parabon NanoLabs (https://snapshot.parabon-nanolabs.com/). Unfortunately, due to poor preservation of all TK individual’s skeletal remains, the standard skull-based facial reconstruction was not possible at the time of this study. Therefore, to illustrate our analysis beyond the basic phenotypic features and to comprehensively reconstruct the facial characteristics of the studied individuals, we used Snapshot® facial prediction based on the imputed calls, which, again, showed the phenotypic diversity indistinguishable from the present-day Koreans (Fig. 6, Fig. S19).

## Discussion

Several recent studies reported genomic results for ancient individuals from China, Japan, Mongolia, Philippines, and Vietnam (*1*–*4*, *6*, *8*, *42*). As the Korean peninsula is located between Japan and China, ancient genomes and population composition from the Korean Peninsula are particularly interesting in the context of historical and archeological findings, including evidence of extensive networks of exchange and contacts between Korea, Northeast Asia, and Japan.

The genetic diversity seen among the Gimhae TK period individuals can be explained by at least two scenarios. One is that the Jomon ancestry observed in Korea-TK_2 is the result of a regional admixture that occurred gradually within the Korean peninsula between at least two subpopulations where the indigenous population already carried the genomic components prominent among Jomon people. There is indeed some evidence of Jomon existence in the Korean peninsula along the Southeastern coastline supported by the artefacts such as pottery and obsidian arrowheads. The obsidian is recognized as an import from Kyushu in Japan (*43*), however such evidence dates to the Neolithic (10000-1500 BCE), a rather distant history. Moreover, Jomon has not been identified as a dominant cultural component in this area, which could imply that individuals bearing Jomon-related ancestry were either completely absorbed by populations originating from northern China, who entered along the Korean peninsula coastlines, or their genetic signal was diluted by various other demographic scenarios, leading to the genetic homogeneity observed in present-day Koreans. Under this scenario, it is possible that the reported DATES result of an admixture event eight generations before the current samples, also can be associated with the admixtures that lead to this dilution. The second scenario is that the population of the Gaya region was diverse and some level of cosmopolitanism prevailed in relation to intense trade and connections with nowadays Japan, similar recently reported results for Iron Age and Antiquity period populations in the case of Rome and neighboring port cities (23). Human movements and admixture are closely related to trading networks and connections between geographic populations in these regions. Archeological studies suggest that the Japanese Yayoi culture received a populational input from the Korean peninsula in a form of populations who crossed the narrow Korean Strait between Busan/Gimhae and Kyushu/Honshu islands (*44–46*) which in some cases may complicate the genetic and cultural differentiation between ancient Koreans and Japanese. Taking into consideration the limitations of the current data, it is difficult to conclusively determine whether the Jomon component in Gimhae samples is derived mainly from local Korean or external Japanese sources. Based on the DATES analysis for the Korea-TK_2 individuals, the admixture with the Jomon component could have occurred around 200 CE, which overlaps with the reign of Gaya, if we assume that there were distinct Jomon-like and non-Jomon-like populations in the region. Moreover, it is important to stress that “Jomon” refers originally to a type of pottery technology and style and is not well defined as a chronological period. There is archeological evidence of multiple subgroups within the broadly referred Jomon people rather than a single homogeneous population. This partially explains why AKG_10203 can only be modelled with the Jomon genome from Ikawazu (897-803 cal BC) (*5*) but not with the genomes from Rokutsu shell Mound (2472-835 cal BC) (*47*). The differences between the dates and geographical locations (Ikawazu genome is from Honshu, Japan’s main island, which is geographically associated with Korea along the well-established Setonaikai coastal route) also highlights the need to consider possible Jomon heterogeneity when interpreting admixture. This however, could also be the result of technical limitations of the method itself due to the low sample numbers.

The use of diploid genotyping and deep-genome sequencing enabled us to identify fine relationships between the ancient Koreans and their present-day descendants. We demonstrated that the people of the TK period in Gaya were more genetically diverse than present-day Koreans, and that they had diverse ancestral components and population substructure that have been homogenizing from the TK period onwards. The phenotypic analysis confirmed that over 100 forensically-relevant SNPs of present-day Koreans were already established during the TK period. For example, *EDAR* V370A (rs3827760), (*36*) the East Asian hair thickness mutation, and the *ABCC11* G538A (rs17822931), the dry earwax variant (*33*), both of which were homozygous in majority of our TK period samples. The same mutations were already reported for Tianyuan man and suggest genetic continuity of this variant in East Asia for the last 40,000 years (*48*). Furthermore, TK period Koreans might have had elevated frequencies of SNPs coding for risk variants of Myopia, which is widespread in Korea (*38*).

Results also suggest that the main burials and the human sacrifices were not associated with separate genetic substructures as both Korea-TK_1 and Korea-TK_2 groups are not based on social status. However, the information inferred from the artefacts was limited due to poor grave preservation and missing artefacts due to tomb robbery acts prior to excavations. For example, male AKG_10203 was identified as a possible main burial of a warrior class or a lower class noble while female AKG_10207 was possibly a human sacrifice. The remaining gilt bronze artefact and arrow quiver ornaments are insufficient to accurately determine the social status of AKG_10203. Due to aforementioned issues, we could not identify the owner of each grave, which is also the case for AKG_10207, who was a sacrifice for a possibly high-class noble as indicated by iron spear and bronze mirror findings. Among the Korea-TK_2 group, AKG_10218 was the main burial while the rest of the samples were sacrifices for different main burials. It is also notable that we could not ascertain the relationship between the sex and social status in Daeseong-dong tumuli, even though the two main burials were of males, it might have been affected by the small sample bias, as human sacrifices clearly belonged to both sexes.

Our findings and the genetic connections between Daeseong-dong people and Jomon-like people calls for future work involving extensive sampling of ancient individuals from South Korea and neighboring regions. Moreover, future studies which focus on intra-cemetery and regional patterns both in East Asia and other world regions are needed in order to develop a more comprehensive perspective about the emergence and diversification of social status and cosmopolitanism among past societies,

## Methods

### Samples

All individuals were sampled with the permission of the local authorities at the Gimhae National Museum, following preventive measures to avoid contamination. Six teeth were collected and twenty petrous bones were drilled using the protocol described in Sirak et al. (*49*). Teeth roots were cleaned and cut using a sandblaster and later pulverized using a mixer mill (Retsch Mixer Mill 400) (*50*).

### DNA extraction, library preparation, and sequencing

All experimental work was performed in the dedicated ancient DNA laboratory of the University of Vienna. Personnel and facilities followed all the standard measures to guarantee the authenticity of the data. We included extraction of negative controls (no powder), library, and PCR negative controls (extract was supplemented with water) in every batch of samples processed and carried them through the entire wet laboratory processing to test for reagent contamination. DNA was extracted from ∼75 mg of powder using an optimized DNA extraction protocol (*50*). Illumina sequencing libraries were constructed using 12.5-25uL of extract and amplified using Accuprime Pfx Supermix (Thermo Fisher Scientific), following Gamba et al. (*51*); a protocol adapted from (*52*). Quality assessment of the amplified library was performed on an Agilent 2100 Bioanalyzer and a Qubit 2.0 Fluorometer. All the amplified libraries were initially screened using NovaSeq platform and further sequenced with NovaSeq platform in the Vienna Biocentre.

### Bioinformatic analysis

Sequencing reads were trimmed using cutadapt (Version. 1.2.1) (*53*) and aligned to the human reference genome (GRCh37), with the revised Cambridge reference sequence (rCRS) mitochondrial genome using BWA (*54*) (Version 0.7.5). Duplicate mapped reads were removed using Picard Tools (*55*). Reads with mapping qualities below 30 were also removed. Unique and filtered reads were analyzed with qualimap-2 (*56*) to assess the coverage of the genomes. MapDamage-2 (*57*) was used to estimate the level of deamination and the authenticity of the data. Reads used for the genotype-calling were trimmed again, 5 nucleotides in both ends, to minimize the genotyping errors in the downstream analyses.

Pseudo-haplotypes were called with a pileup caller of sequenceTools (*58*), filtering out bases with base-quality score below 30. In this stage, we called the positions of the 1240k dataset (*25*). These calls were merged with individuals from (*2*, *4*, *47*, *59*–*63*) and the rest of present-day genomes available from David Reich group’s website (reich.hms.harvard.edu). The final dataset is composed of 645 present-day Eurasian (*47, 64*) samples and 237 ancient individuals.

### Data authenticity

Contamination level was estimated from the mtDNA sequences using Schmutzi(*27*) and by estimating the heterozygous content of the male X chromosomes using ANGSD (*65*). Mapping qualities and read length distribution were also examined with FASTQC (*66*), the average genomic and mitochondrial depths were obtained using Qualimap (*56*).

### Uniparental Markers

Reads aligned to mitochondria were processed with Schmutzi (*27*) to generate a consensus sequence of the mitochondrial genomes. We called the mitochondrial haplogroups of the mitochondrial consensus sequences using haplogrep 2.0 (*28*) and the version 17 of Phylotree. Y chromosome haplogroups were estimated using Yleaf (*67*).

We used lcmlkin 1.0.0 (*68*) to explore the presence of kinship relationships between the samples analyzed.

### Population Genetic tools

All the subsequent analyses were performed with the 1,240k dataset.

Principal component analysis was conducted using 597,573 SNPs and 645 present-day genomes using smartpca from Eigensoft package (*69*). Resulting data was plotted using R (*70*). Ancient samples were projected into the PCA that was based on the present-day ones, using option “lsqproject”. Two rounds of outlier removal were used. Results with ancient samples were plotted using R.

An unsupervised ADMIXTURE analysis was performed with ADMIXTURE 1.3.0 (*71*) with the modern genotypes from the publicly available Human Origins (HO) panel (*69*) together with the eight 1,800 year old Korean SNPs, restricting the analysis to the 597,573 SNPs of the HO dataset (*64*). These SNPs were filtered for MAF < 0,05 and Missing sites > 0,05. Filtered SNPs were pruned for linkage-disequilibrium (LD) using PLINK 1.9 (*72*) flag --indep-pairwise with a windows size of 200 SNPs, advanced by 50 SNPs and establishing an r2 threshold of 0.4. A total of 282,896 SNPs were used in the analysis. The final ADMIXTURE was run with K ranging from 2 to 15 and 100 bootstrap replications. The ADMIXTURE result was plotted with PONG 1.4.9 and in R (*73*).

For the analysis of the differential X chromosome ancestry, we run ADMIXTURE with a pruned dataset of 3,063 SNPs and samples that exhibited less than 90% of missing sites. This has resulted in 56 samples out of the 237 ancient genomes included in the initial dataset. The analysis has been performed and analyzed as general ADMIXTURE analysis described above.

*f*_3_-statistics were run using admixtools (*69*) in the form *f*_3_(X, Test; Mbuti) using all the populations of the dataset, both present-day and ancient. *f*_4_-statistics were also run using the same package. We used the form *f*_4_(X, Test; PopA, PopB) using all the possible combinations of a subset of 31 populations (Table S7). Only results with |Z| > 3, and supported by more than 100,000 SNPs were considered.

### qpWave and qpAdm

We used the software *qpWave* (*74*) from ADMIXTOOLS 5.1 (*69*) with parameter “allsnps: NO” to investigate the homogeneity of the ancient individuals and if any of them would require additional sources of ancestry to explain their genomes. In order to capture a wide range of distal ancestries we used the following base “Right” outgroup set of populations (“R10”): Mbuti.DG, Ami.DG, ONG.SG, Mixe.DG, Tianyuan, Iran_GanjDareh_N, DevilsCave_N, Bianbian, Liangdao2, Yumin. We used a tolerant threshold of p=0.01.

In admixture modelling we used *qpAdm* (*25*), again with parameter “allsnps: NO”, to find populations and potential admixture events from which the two ancient Korean subgroups could have descended from. As the outgroup “Right” set we used the base R10. As sources we used a set of populations based on the top results from the outgroup-*f_3_* statistics of the form *f_3_*(X, Test, Mbuti), for Korea-TK_1 and Korea-TK_2: Amur_Basin-IA, Ekven_IA, Japan_Jomon, Japan_JomonIK002, LateXiongnu, LateXiongnu_Han, Yankovsky_IA, Yellow_River-BA, Taiwan-IA, Liao_River-BA. We applied a more stringent threshold of significance, when compared with qpWave, of p=0.05. As part of a 2-step model competition approach, models passing this threshold were re-evaluated and only accepted if they remained significant after moving all unused source populations to the “Right”.

### Treemix

We employed Treemix 1.13 (*75*) to estimate the historical relationships among populations, using a graph representation that allows both population splits and migration events. We ran Treemix with five repetitions using from 0 to 8 migration events (-m), with a -k value of 500 and with low coverage correction (-noss). The final list of populations has been selected using the *f_4_* results and restricting the analysis to the populations that did not exhibit high error values.

### Dates

We used DATES 753 (*76*) to estimate the dates of different admixture events, based on the results of qpAdm. We ran the program using parameters as follows: bin size: 0.001, maxdis: 1.0, runmode:1, qbin: 10 and jackknife: YES.

### Diploid Calls and imputation

We first processed the filtered mapped reads with RealignerTargetCreator from GATK 3.7 (*77*), and realigned the indels from the Mills_and_1000G_gold dataset. Afterward, we added MD tags to the bam reads using samtools 1.9 calmd (*54*). SNPs overlapping both the 1,000 genomes dataset and the 1,000 Korean genome project (Korea1K) set (after lifting it over to hg19 coordinates) (*1*) were called in these reads using GATK UnifiedGenotyper, setting the options-out_mode EMIT_ALL_SITES and --genotyping_mode GENOTYPE_GIVEN_ALLELES. Option -glm SNP was also used to get the genotype likelihoods of the SNPs. Obtained calls were filtered to add equal likelihoods to both missing data and the sites possibly affected by deamination. Resulting files were split into chromosomes with bcftools 1.6 (*78*) and filtered for sites with MAF>0.05, and less than 10% of genotype missingness. Resulting files were splitted with splitvcf.jar from Beagle in 100,000 SNPs files, with 20,000 SNPs of overlap. We used Beagle 4.0 (*79*) to impute the genotypes setting “gl” option, “gprobs=true”, “window=15000”, “impute=true” and giving maps and reference files from Beagle. Imputed calls were filtered by Genotype probability > 0.99, MAF > 0,05 and excluding transitions. The resulting datasets consist of 1,574,651 filtered and imputed diploid calls. We repeated the same procedure using the 1,000 genomes data only, in order to obtain a second dataset with all the SNPs covered by 1,000 genomes imputed.

### Chromosome painting analysis (finestructure)

We used vcftools 0.1.13 to merge our eight imputed samples with modern East Asian samples from 1,000 Genome Project (*80*) (Han, Dai, Korean, Japanese, and Kihn_Vietnamese), a Korean set from Jeon et al. (*10*), and six additional ancient shotgun genomes from East Asia (NEO204, NEO236, Kolyma_River_M, M9984 and IK002) (*4, 59*). The total number of SNPs after filtering out all the positions with genotype missingness “--max-missing 1”, Hardy weinberg equilibrium “--hwe 0.000001”, and minor allele frequencies “--maf 0.00000000001”, was 424,255. Relatives were excluded from the present-day populations utilizing KING v.2.2.4 with options “--related” and “--degree 3”. The ethnic representatives for each present-day population were randomly downsampled to leave a) 10; b) 5; c) 3 representatives per each group. The finalized datasets were converted to map and ped formats using plink v1.90 with option “--recode 12” to preserve the phasing information. We ran scripts “plink2chromopainter.pl” and “makeuniformrecfile.pl” from the FineSTRUCTURE pipeline “fs_4.1.1” to generate phasing and recombination information files. Effective population size (*Ne*) was estimated using ChromoPainterv2 with 10 random samples and settings: “-a -i 20 -in -iM”. The average *Ne* was estimated using “neaverage.pl”. We ran FineSTRUCTURE v.4.1.1 in linked mode with options “-x 1000000 -y 1000000 -z 10000” and “-y 10000 -m T”. The results were visualized in FineSTRUCTURE GUI for Windows v.0.1.0 (https://people.maths.bris.ac.uk/~madjl/finestructure/finestructureGUI.html).

### IBD analyses

IBD blocks were identified using a Refined IBD version “17Jan20.102” with default settings and a recombination map provided with the program. For an input, we merged our eight imputed ancient Korean genomes with East Asians from 1,000 Genome Project (*80*), a Korean set from Jeon et al. (*10*) and six additional ancient shotgun genomes from East Asia and filtered them as described in the “Chromosome painting analysis” section above. Our eight imputed samples were generated and prefiltered as described in “Diploid Calls and imputation” section. Additionally, all the sites with missing variants and fixed alleles (MAF>0) had been excluded, yielding 424,255 high-quality markers used as an input.

IBD segments with LOD>3 were analyzed, excluding the few sporadic large-size segments (>39CM) that are most likely an artifact (shown in S19). We averaged the values of segment length and segment counts to reflect the pattern of genetic material shared between each ancient sample and the present-day East Asian populations. The results were plotted using heatmap.2 in R.

### Phenotypic variant analysis

From the imputed ancient genomes, we selected 160 variants with phenotypic implications, retrieving the alleles and their “rs ID” from dbSNP. The calls included in Table S14 correspond to the genotype probabilities imputed with the Korea Genome project and 1,000 genomes overlap-based merged panel (denoted in black font) or the 1,000 genomes alone (denoted in blue font). Genotypes were filtered by genotype probabilities of 0.9. For reference, we included their allelic frequencies in East Asian populations.

## Acknowledgments

We would like to thank Kendra Sirak and Éadaoin Harney for software help and discussions, and Yeonsu Jeon for assitance in aesthetic visualization. We thank Dan Bolser and Whan-Hyuk Choi for giving critical feedback with editing and Jaesu Bhak for editing the manuscript. We are grateful to Gayoung Park (University of Washington), Su-Whan Kim (Gyeongnam Provincial Government Gaya Cultural Heritage Division, Korea), Seong Won Cho (Pukyung National Museum), Weon Young Song (Daeseong-dong Tombs Museum) for help with archaeological context and information.

## Author Contributions

DF and RP collected the samples, JH, JY, YK, SY provided the samples and archeological and historical context. DF, OC, KTO, SS and VO performed the lab work, PG, AB, SJ, HC, and DF analyzed the data. EG organized face predictions for ancient genomes. PG, AB, DF, JB, and RP wrote the paper with input from all co-authors. YC and KB organized sampling and archeological data.

## Data and materials availability

The sequencing data is accessible in the European Nucleotide Archive (ENA) under the accession code: PRJEB45573.

## Supplementary Materials

### Supplementary Text

#### 1. Archeological context

##### 1.1 Daeseong Dong

The archeological site of Daeseong-dong, Gimhae in Gyungsangnamdo province, is situated on the hill facing the South Eastern coastline of South Korea between the Gujibong Peak and Bonghwangdae, and east of the Surowangneung tomb (Tomb of King Suro). The tombs of this site were built during the Geumgwan Gaya period (42-532 AD). The burial complex of Daeseong-dong together with the complex of Bokcheon-Dong are the most well-known sites of the Gaya Royal Class. The funerary complex covers an area of 3700 m^2^. Since fieldwork started in 1990, 219 individuals have been recovered (*81*), of which 69 are buried in large tombs made of wooden coffins, that believed to be the ones belonging to the ruling classes (tomb owners from now on). Most of these burials include human sacrifices and grave goods.

During the Three Kingdoms period, Gimhae was a major trading port, as is evident from the multiple Chinese and Japanese artefacts recovered from the tombs (Figure S1). One example is the mortuary finds from tomb 91, dating to the early 4^th^ century AD which include a bronze vessel made in Northeast China, a gilt-bronze harness made from the Conidae shells that was imported from Japan, bronze horse bells, a chamfron, and cylindrical bronze implements from Japan. Substantial Japanese bronze ornaments were also unearthed from tomb 88 that dates to the mid 4^th^ century AD.

Iron artifacts, including armors, iron helmets, and coins, were very common in most tombs. One of the most remarkable finding were the grave goods from tomb 29. The owner of tomb 29 was buried inside a large wooden coffin built in the mid 3^th^ century AD, that lies over more than 100 iron ingots. Additionally, this tomb consisted of multiple pieces of pottery and bronze pots which are stylistically typical of a Northern-style culture. This culture was introduced to the Gimhae area at the end of the 3^rd^ century AD, probably from the Northern nomadic steppe tribes (*82, 83*). The high status burials of this culture are characterized by the presence of Gimhae-type wooden-chambered burials, Dojil pottery, Ordos-type bronze pots, iron armors and trappings for horse riding, typical of nomadic tribes. This has been understood as a signal of intensive trade between the northern East Asian region and Gimhae area. But it is unclear as to whether this could also indicate the migration of northern tribes, especially from the Buyeo Tribe (An ancient Korean Kingdom located in Middle Manchuria) (*84*), as no anthropological or genetic studies have been carried out to test this hypothesis. The appearance of these non-local objects also corresponds to the timing of the appearance of human sacrifices in the complex, at the end of the 3^nd^ century AD and to the burial division starting from the 4^nd^ to 5^nd^ centuries AD as indicated by the appearance of Gimhae-type wooden-chambered burials. The majority of burials and objects associated with ruling class tombs are from the 4^th^ and 5^th^ centuries. The tombs of the rulers are placed on the upper parts of the hills (*85*).

The ritual of human sacrifice as a funeral practice first appeared in the Gyeongsang province, and especially in Gimhae. No evidence of sacrificial rituals was reported from other Korean regions. The first human sacrifices appeared in the Gimhae region at the end of the 3^th^ century, and became prominent during the 4^th^ century, and then rapidly disappeared in the Gimhae area during the late 5^th^ century. The sacrificed persons in Daeseong-dong Tumuli were previously considered as slaves, but recent studies suggest them as warehouse keepers, concubines, horsemen, and guards related to the main burial. It is not possible to discern the relationship between the owner of a tomb and sacrifice victims. Anthropological data suggest that human sacrifices in Gaya were mostly in their 20s and 40s. Women sacrifices are a bit more common than men and this pattern could be related to the kind of their labor (*82*, *86*, *87*).

##### 1.1.1 Individuals included in the study

###### A) Individual AKG_3421, burial 11

Individual AKG_3421 (Ancient Korean Gimhae Genome 3421) is a female sacrifice found in tomb 11. Individual AKG_3421 is found together with the tomb owner of the burial. The main buried individual is found in a single wooden coffin. The tomb is indirectly dated to the early 5^th^ century AD based on the coffin characteristics.

###### B) Individual AKG_10207, burial 23

Burial 23 is a single wooden coffin with four human sacrifices and the burial is dated to the late 4^th^ century AD, based on the burial style. Individual AKG_10207 is female sacrifice B of this burial, a non-privileged individual. Japanese and Chinese style relics were discovered in the tomb. The funerary objects; bronze mirror and iron spear, suggest that the burial owner belongs to a high-class individual.

###### C) Individual AKG_10203, Burial 12

Individual AKG_10203 is the owner of tomb. He is found inside a wooden coffin together with gold and bronze artifacts from the early 5^th^ century. Probably this individual belongs to a warrior class or to the low-nobility, as the tomb is not one of the richest ones. He is therefore classified as a privileged individual.

###### D) Individual AKG_10204, Burial 12

Male sacrifice of tomb 12, representing a non-privileged individual.

###### E) Individual AKG_10209, Burial 24

Individual AKG_10209 is the sacrifice B of Burial 24, of a non-privileged female. The owner of Burial 24 is buried in a wooden coffin together with three sacrifices dating to the early 5^th^ century AD. One spindle was found in the burial, which is a rare object for the site. The owner of the burial is a privileged individual.

###### F) Individual AKG_10210, Burial 24

Individual AKG_12010 is the Sacrifice C of Burial 24.

###### G) Individual AKG_10218, Burial 62

Individual AKG_10218 is the tomb owner of Burial 62, the relics found in the burial date to the mid 4^th^ century AD. The objects found in the tomb suggest a privileged individual. This individual is a male.

##### 1.2 Yuha-Ri, Gimhae

The Yuha-ri site, also known as Monument Gyeongsangnam-do No. 45, is located on a slope about 15m above the sea level in the city of Gimhae. (Figure S2). Multiple artefacts have been documented from this site, including extensive Joseon Dynasty artefacts found in Trenches 1 and 2 that include porcelain, pottery, and tiles.

Trench 3 is 17 m long and 2 m wide and 60 cm deep trench along the southern walls of the complex. A shel-rich layer at the eastern section of the trench, at a depth of 20 cm yielded (in addition to the numerous shells) Three Kingdoms period vessels and soft earthenware. A five years old baby (individual AKG_3420) was excavated at the western part of the trench; a Joseon dynasty tomb was identified about 30cm below the topsoil (Figure S3). The ceramics and other artefacts that were unearthed surrounding the body indicate that the burial dates to 4^th^ century AD (Three Kingdoms Period). No other tombs were found on this site(*88*).

An anthropological study was carried out in 2017 in the Bioanthropology Lab (Department of Anthropology, Seoul National University) using standardized anthropological methods (*89*). Based on the length of the long bones the age estimate is 3-6 years old (*90*). All of the deciduous teeth have erupted and the dental analysis suggests that the child is at least 3 years old (±1 year old). Considering the growth status of the crowns of the permanent dentition the subject is estimated to be 4 years old (±1 year old).

#### 2. Extended Methods Diploid calls analyses

We realigned the indels in our bam files with GATK 3.7 (*77*) IndelRealigner. We also trimmed two bases per read end to minimize the effect of damage in the variant calling using trimbam from bamUtil 1.0.14 (*91*). GATK Unified Genotyper was used to call variants using the known alleles from 1000 Genomes (--alleles) and restricting the calling to SNP with genotype likelihoods (-glm), we used -out_mode EMIT_ALL_SITES. Called variants were later filtered with bcftools 1.12 (*92*), excluding transition sites. The resulting genotypes were also filtered excluding sites with more than 10% of missing sites.

Resulting sites were split by chromosomes with vcftools 0.1.16 (*93*) and we used the script splitvcf.jar to split the VCF files in files of 100,000 with an overlap of 25,000. These genotypes were imputed with Beagle 4.0 (r1399) (*94*). Using the genetic maps provided by Beagle. As a reference panel we used the genotypes of Asian individuals from 1,000 genomes and the individuals from the 1,000K genome project. Afterwards we filtered the sites for GP>0.99, MAF 0.05 and excluding transitions. The resulting number of sites is 1,574,651 SNPs.

---

**Fig S1:**
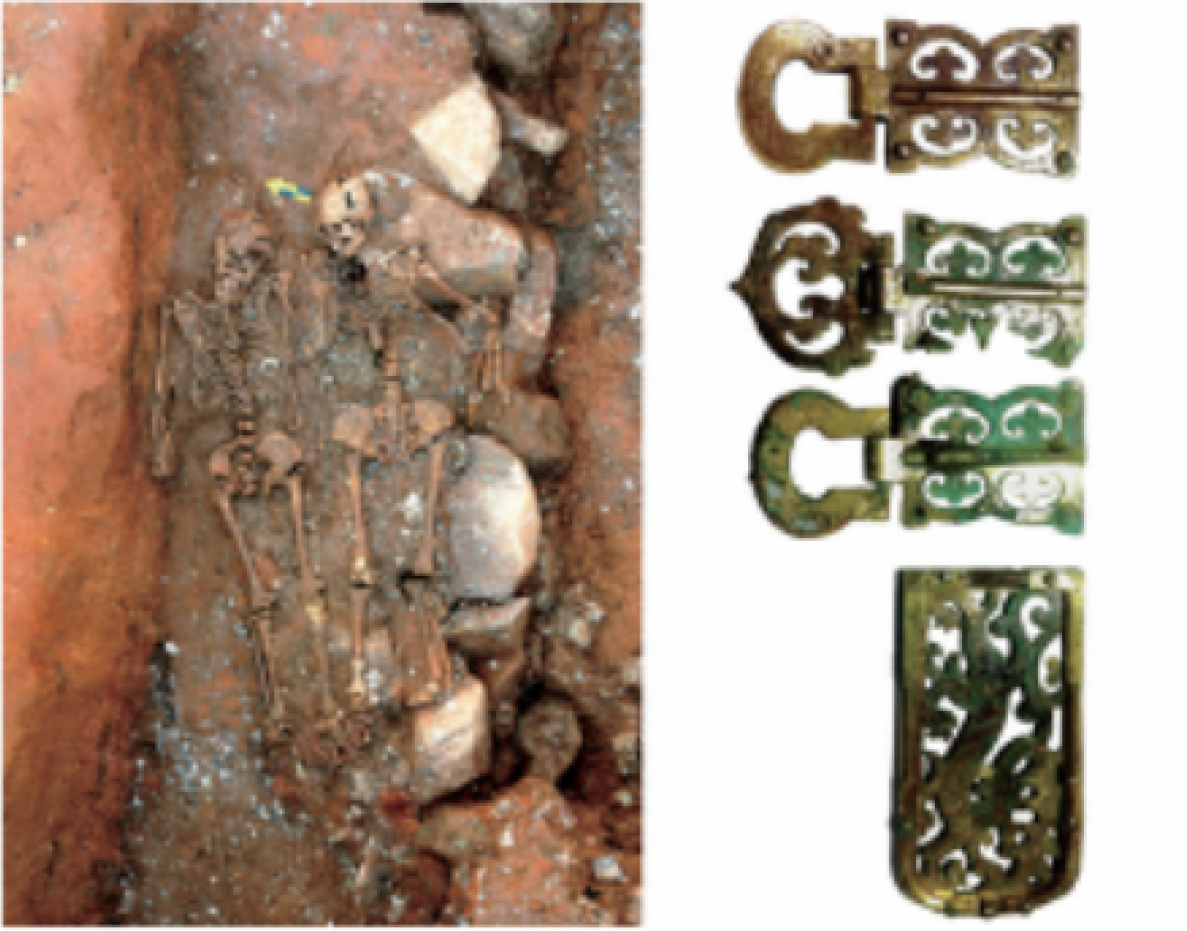
Human sacrifices and Japanese style funerary bronze ornaments found in tomb 88 of Daeseong-dong complex.

**Figure S2:**
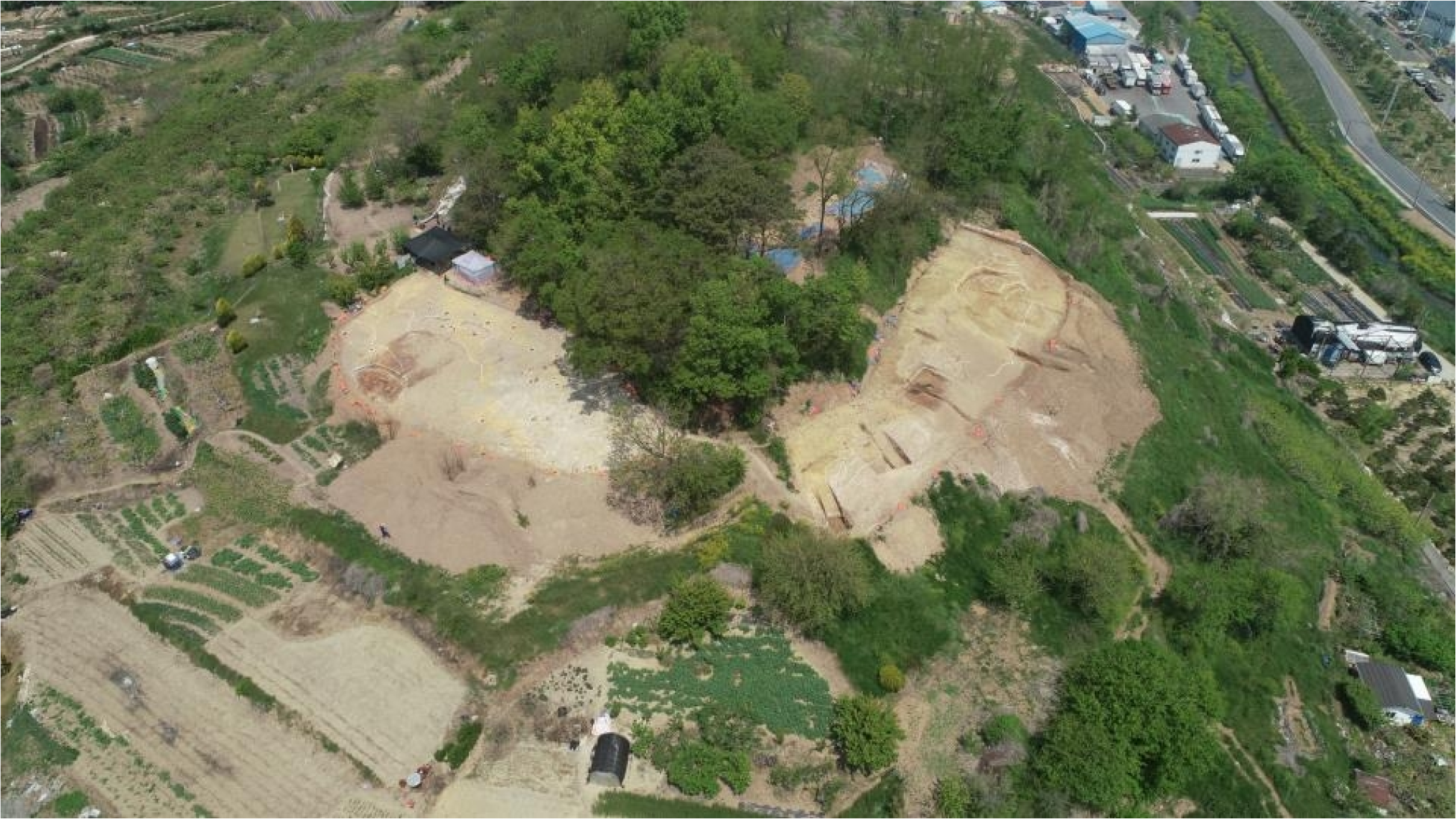
General view of Yuha-ri site in Gimhae.

**Figure S3:**
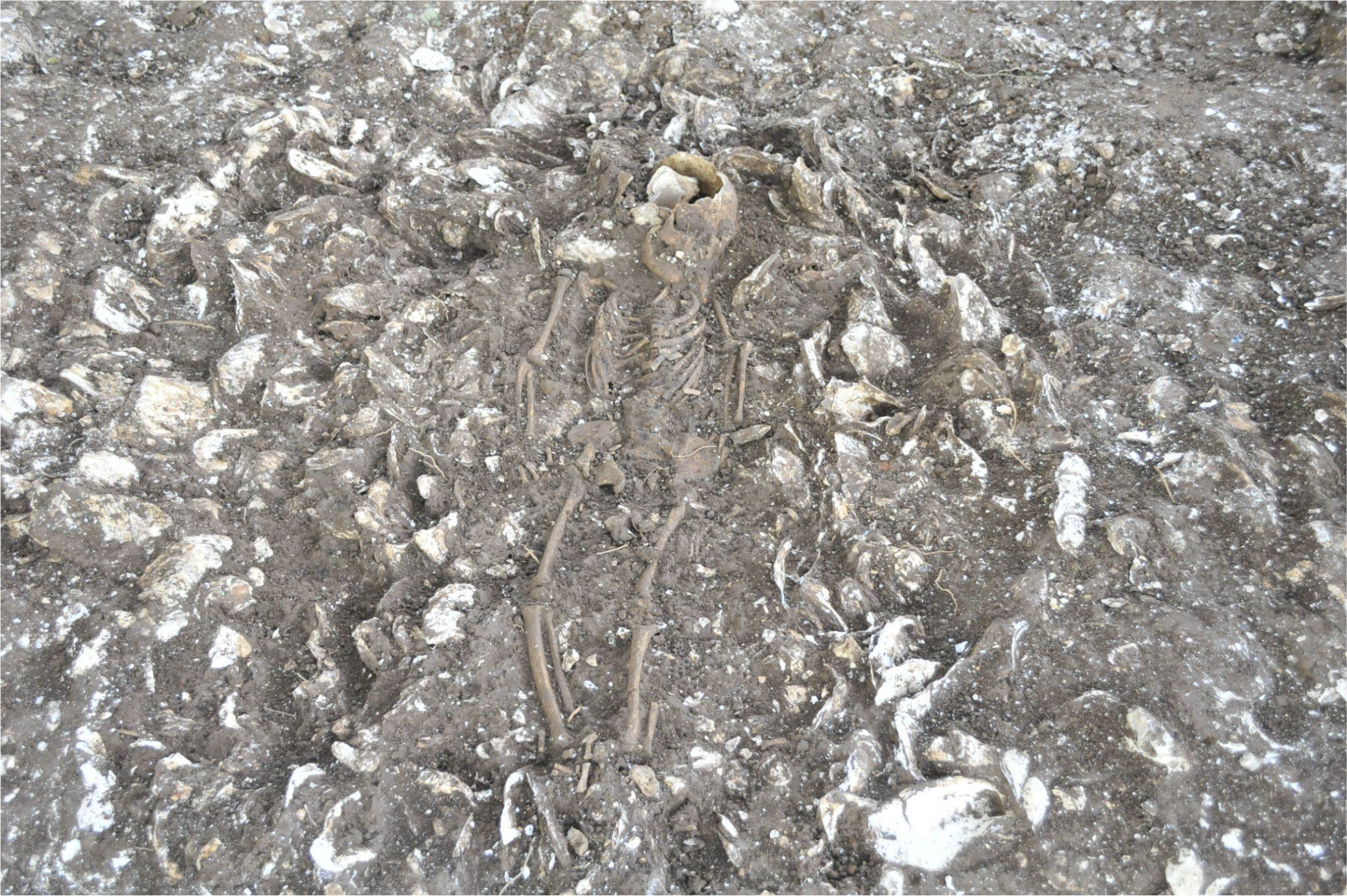
Image of the burial of individual AKG_3420 in Trench 3.

**Fig. S4:**
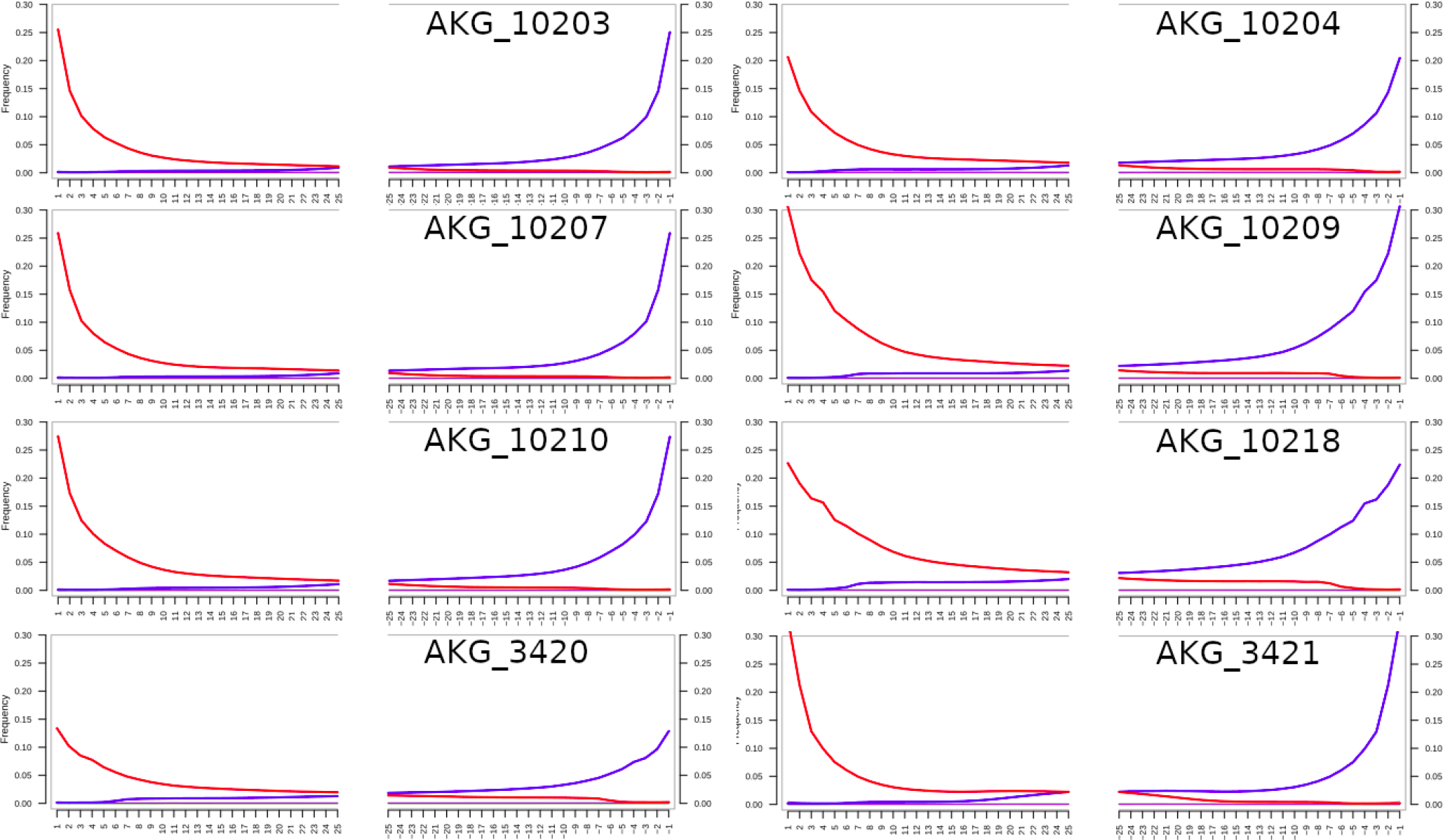
Deamination profiles of the eight ancient Korean genomes. All the samples show the typical patterns of ancient DNA deamination.

**Fig. S5:**
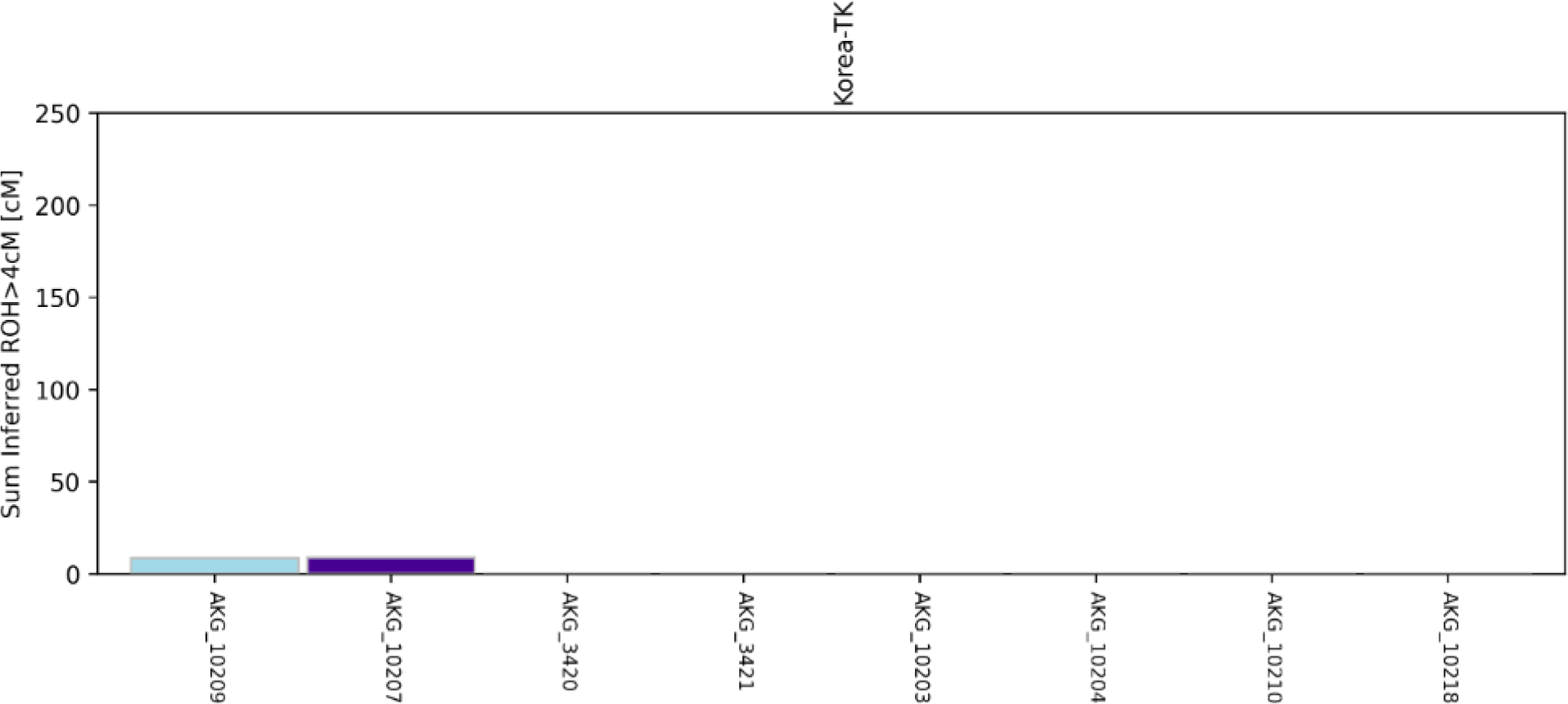
ROH results of the Korean individuals. Y axis shows the sum of ROH longer than 4 cM. Light blue shows ROH between four and eight cM and blue shows ROH between eight and twelve cM. It is observable that the studied individuals show almost no RoH in their genome.

**Figure S6:**
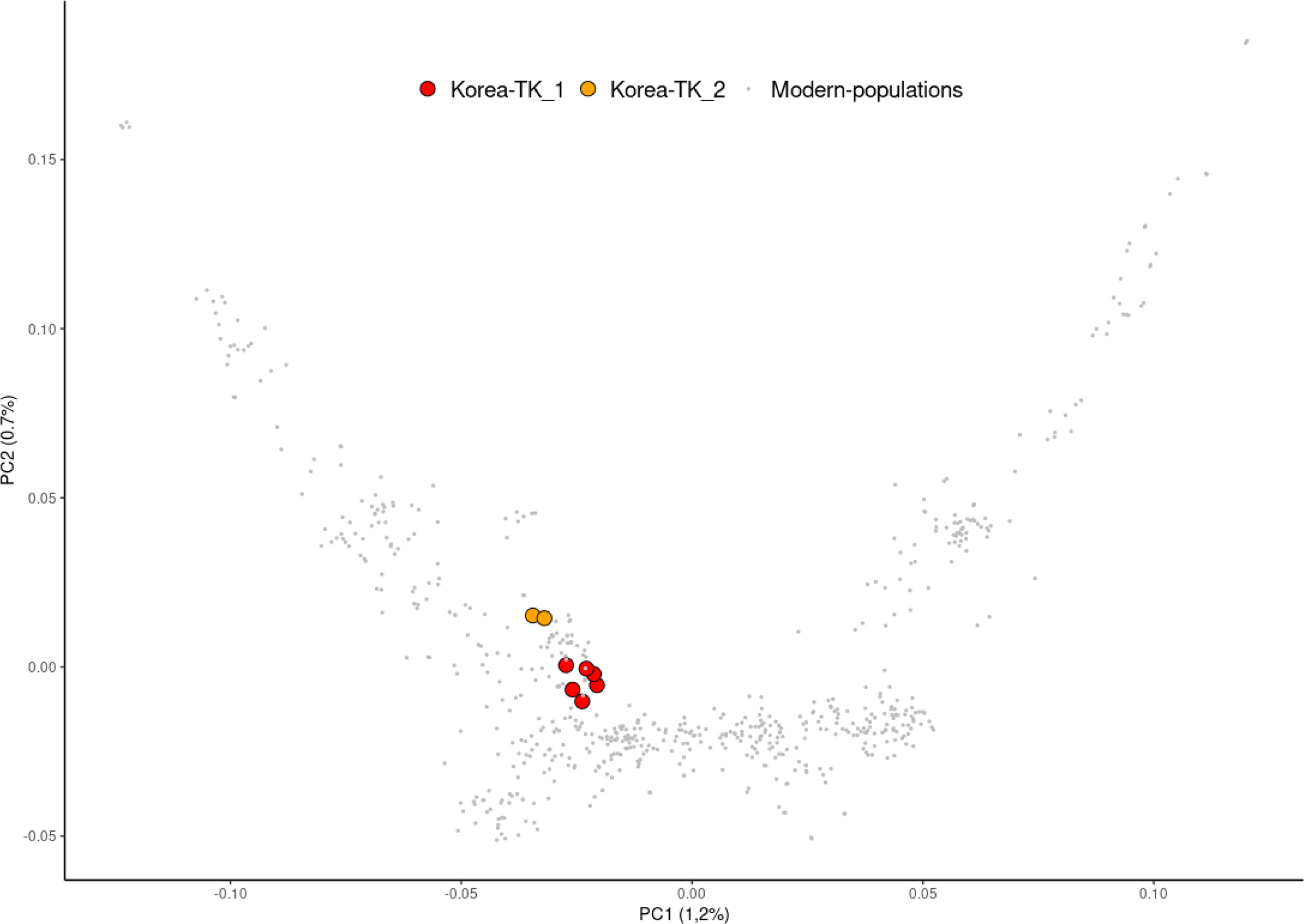
PCA analysis built with the modern samples and the 8 ancient Korean genomes projected.

**Figure S7:**
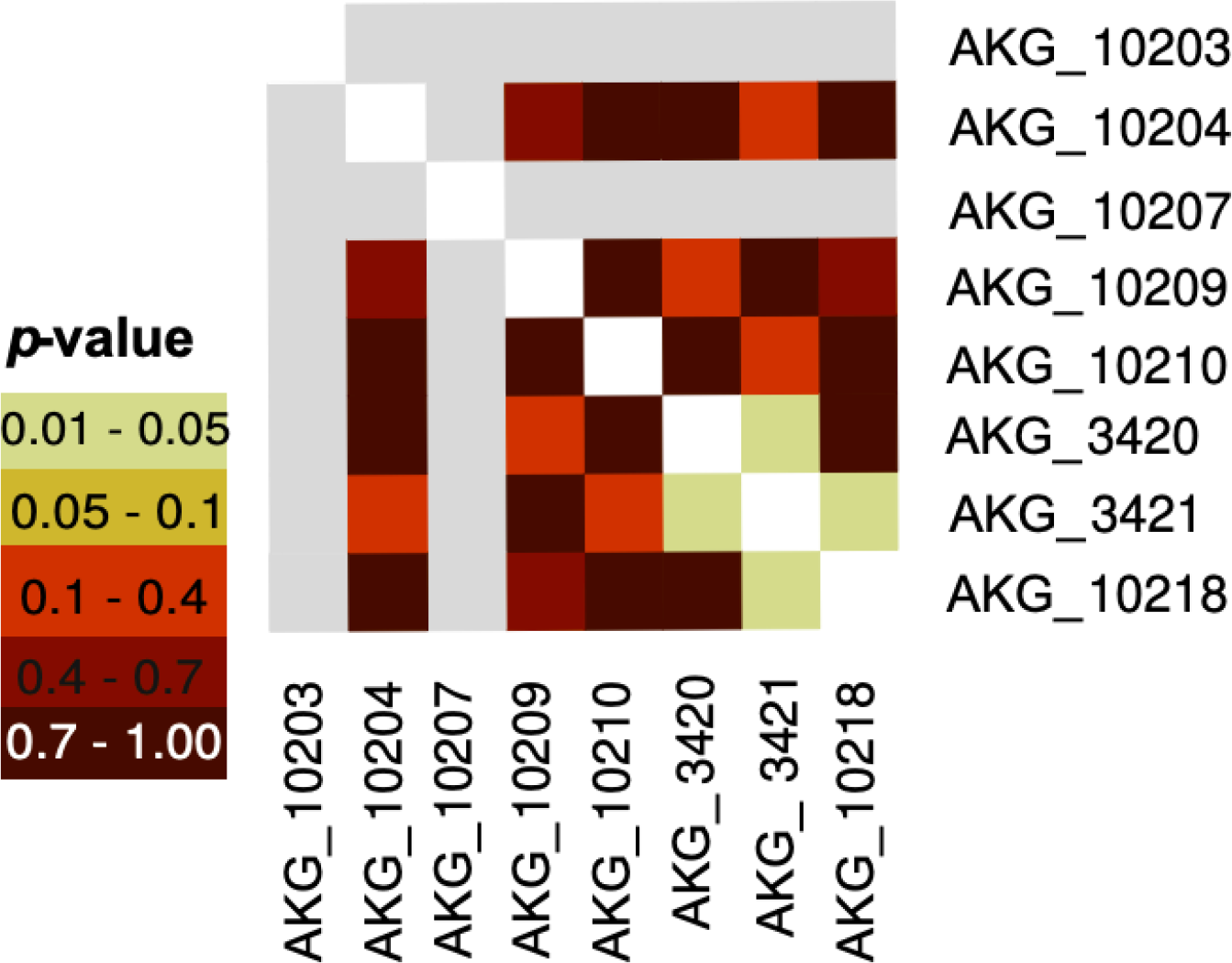
qpWave plot: the qpWave analysis shows the presence of two differentiated clusters. *f_4_*-statistics: Korea-TK_1 is associated with ancient populations from Northern China, while Korea-TK_2 has the closest affinity with Amur Basin and Japan_Jomon populations.

**Figure S8:**
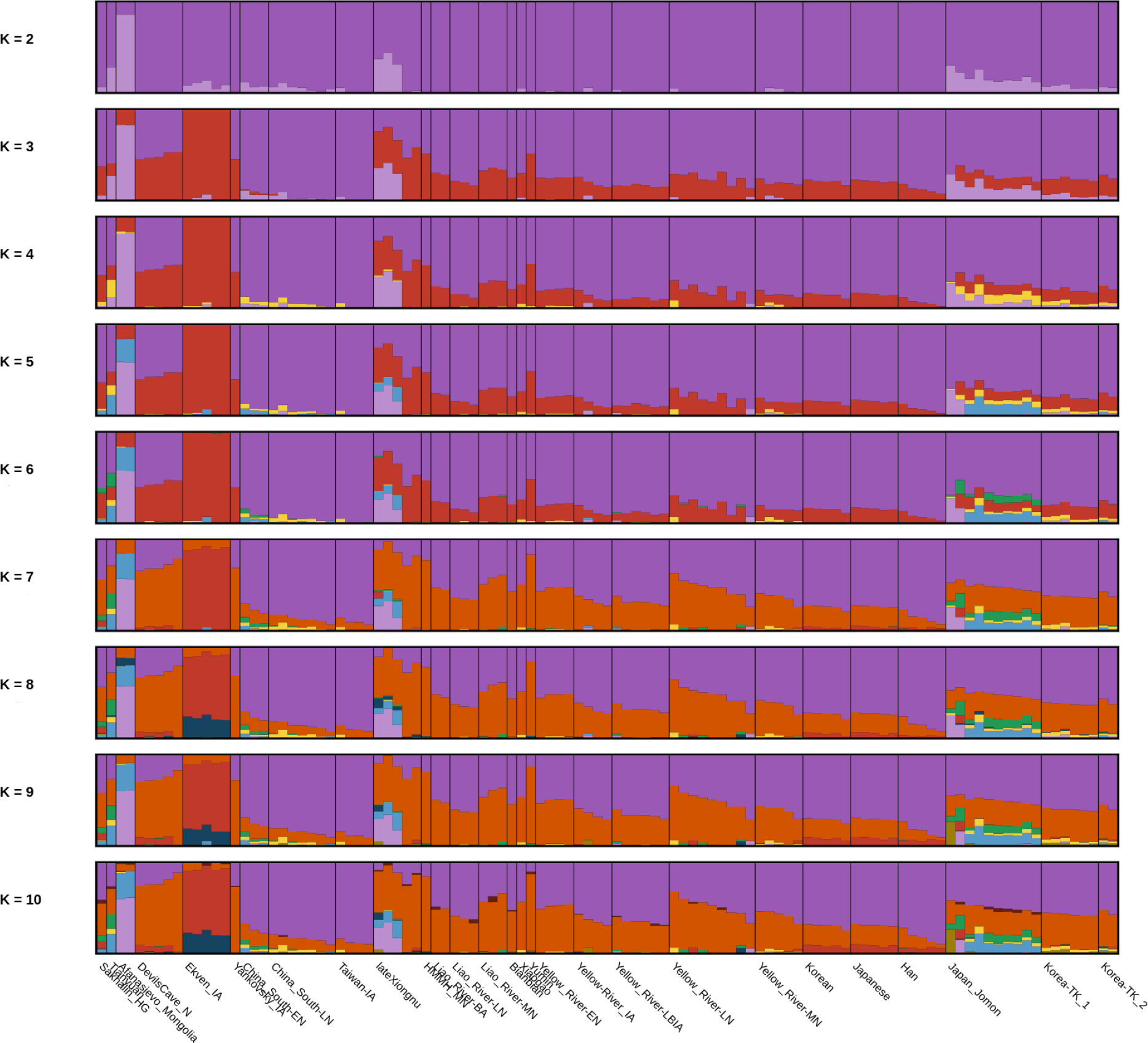
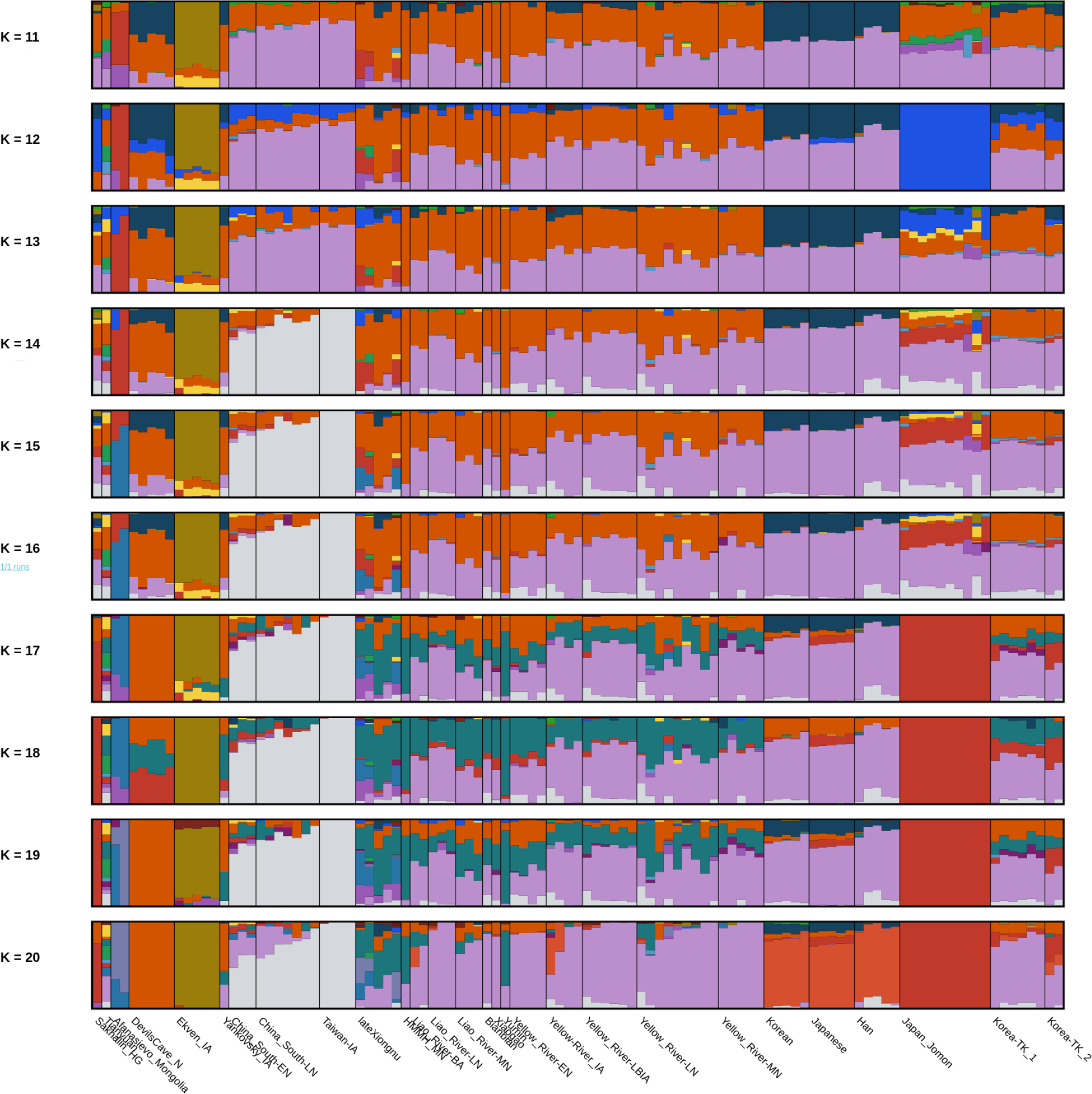
Admixture plots from K2 to K20. We have plotted 108 selected representative individuals to illustrate the genetic composition of the Three Kingdoms period Gimhae Korean genomes.

**Figure S9:**
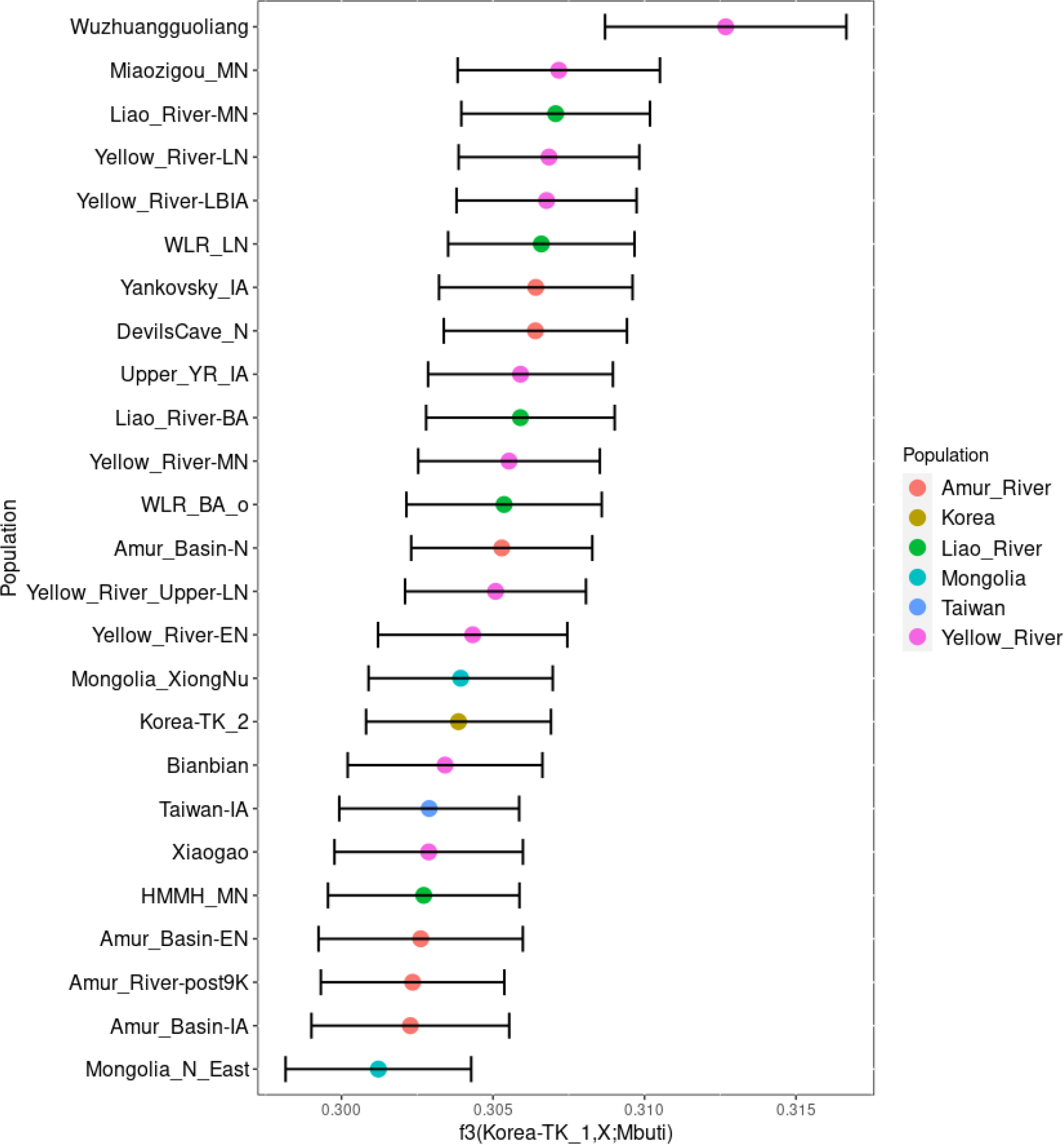
*f_3_*-statistics analysis on the comparison *f_3_*(Korea-TK_1,X;Mbuti). The plot includes the populations with the highest *f_3_* values with at least 50,000 SNPs. The error bars show 1 SD. The colors indicate the population clusters.

**Figure S10:**
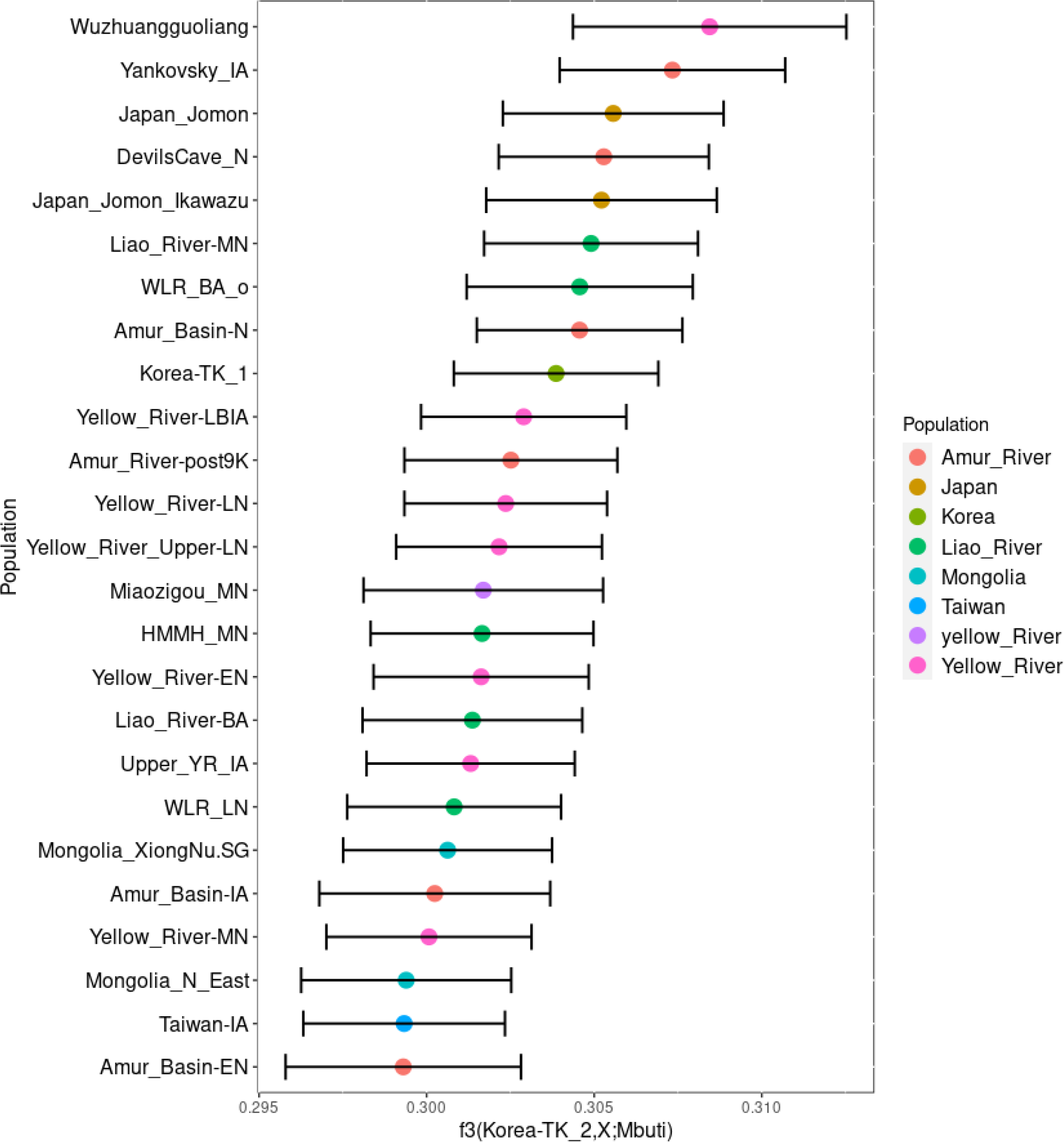
*f_3_*-statistics analysis on the comparison *f_3_*(Korea-TK_2,X;Mbuti). In the plot, populations are shown with *f_3_* values using at least 50,000 SNPs. The error bars show 1 SD. The colors indicate the population clusters.

**Figure S11:**
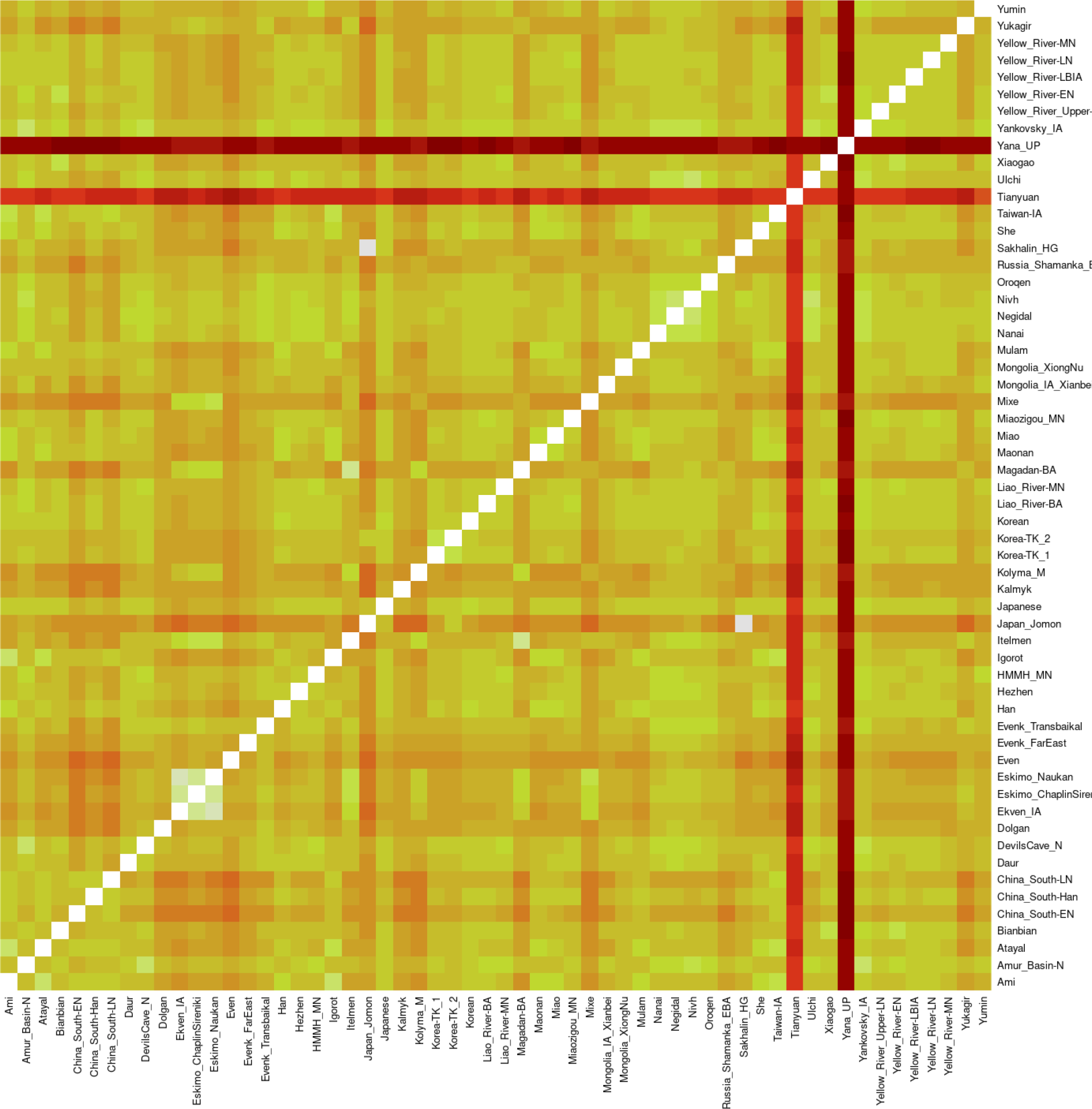
Heatmap analysis on the *f_3_* comparison with all the possible combinations. The plot is built with the *f_3_* values with at least 50,000 SNPs.

**Figure S12:**
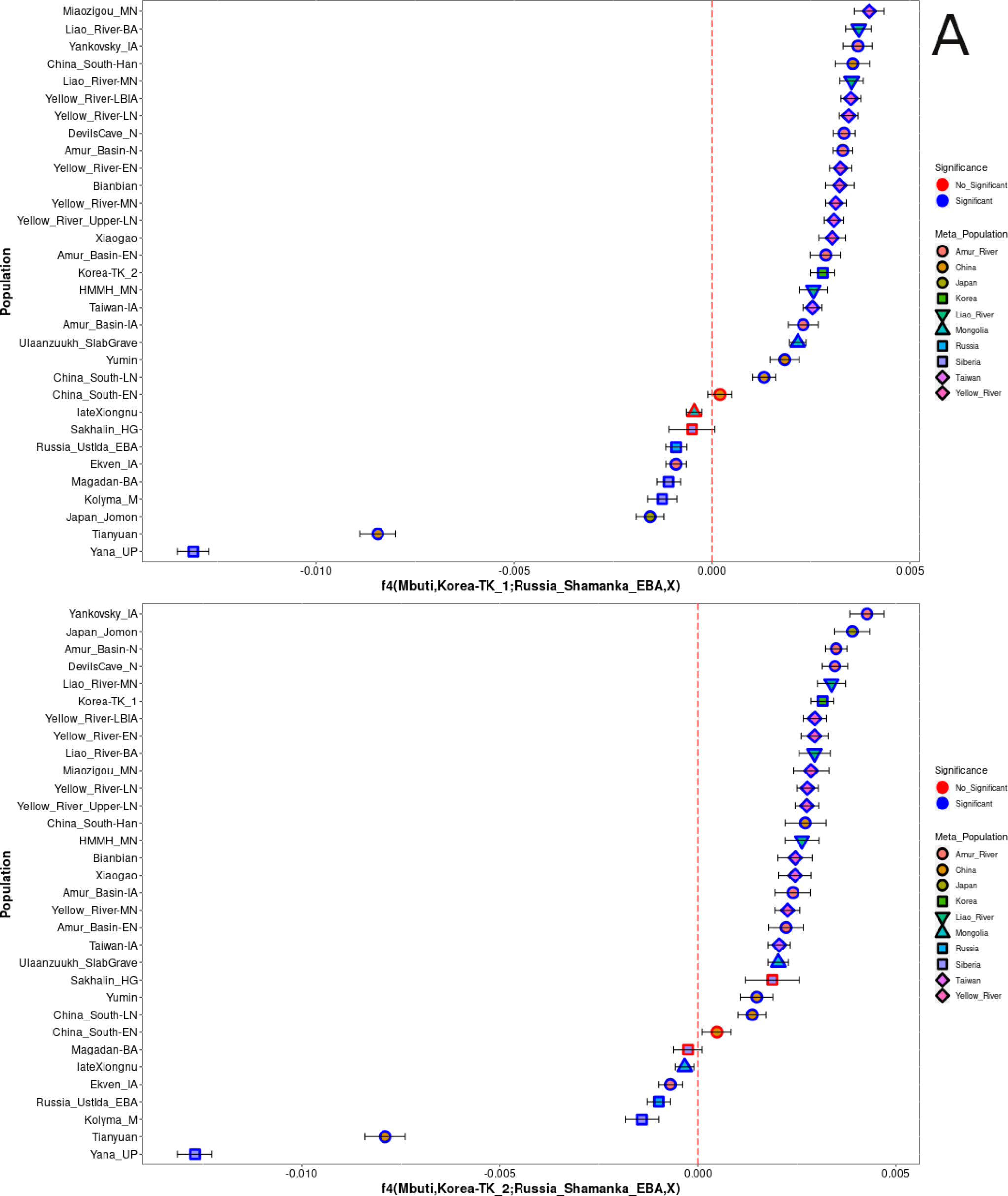
*f*_4_ comparison analysis using (Mbuti,Korea;Russia_Shamanka_EBA,X)

**Fig. S13:**
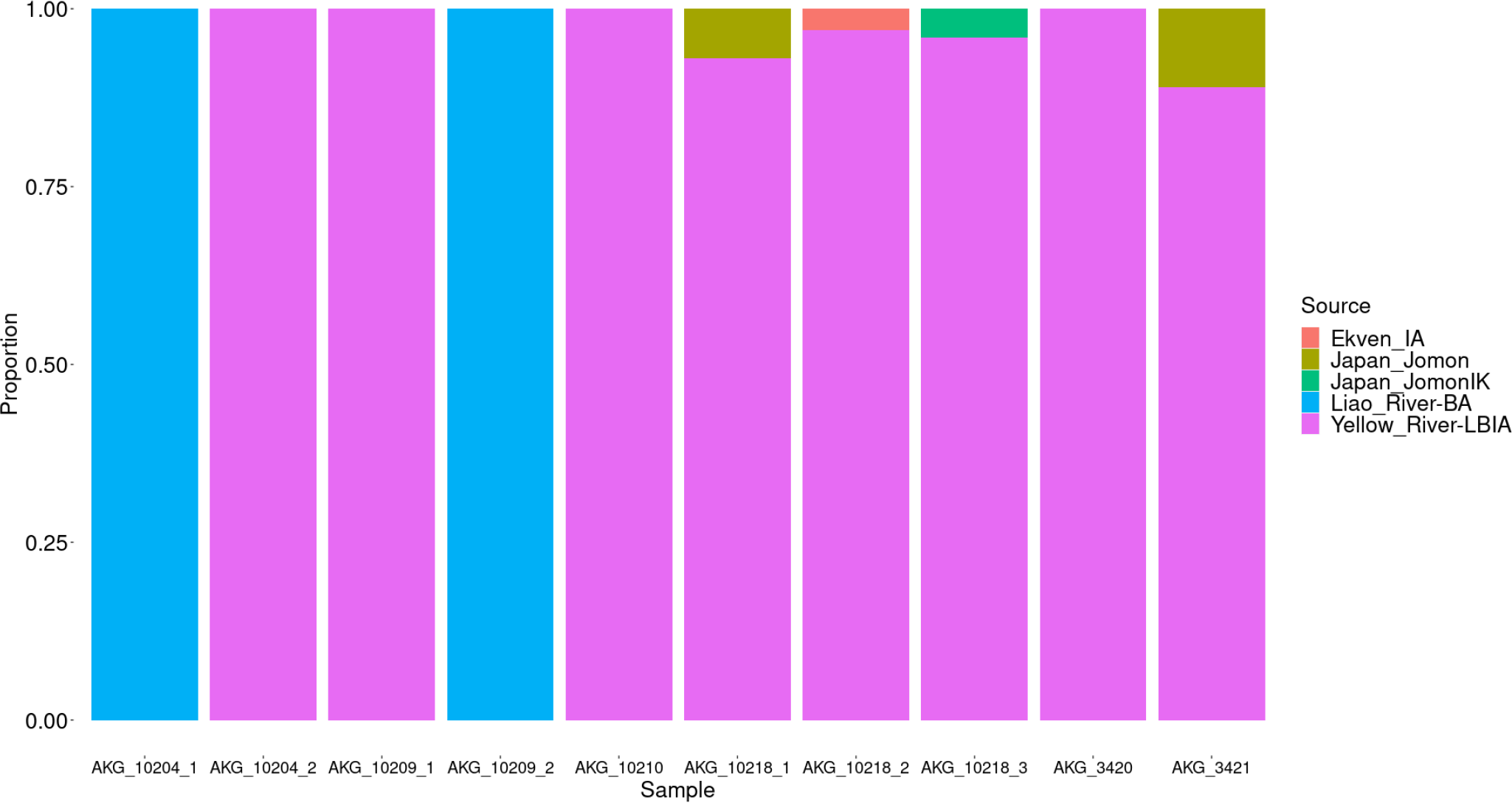
qpAdm results of all the working models of Korea-TK_1. The appendix numbers indicate the multiple models.

**Fig. S14:**
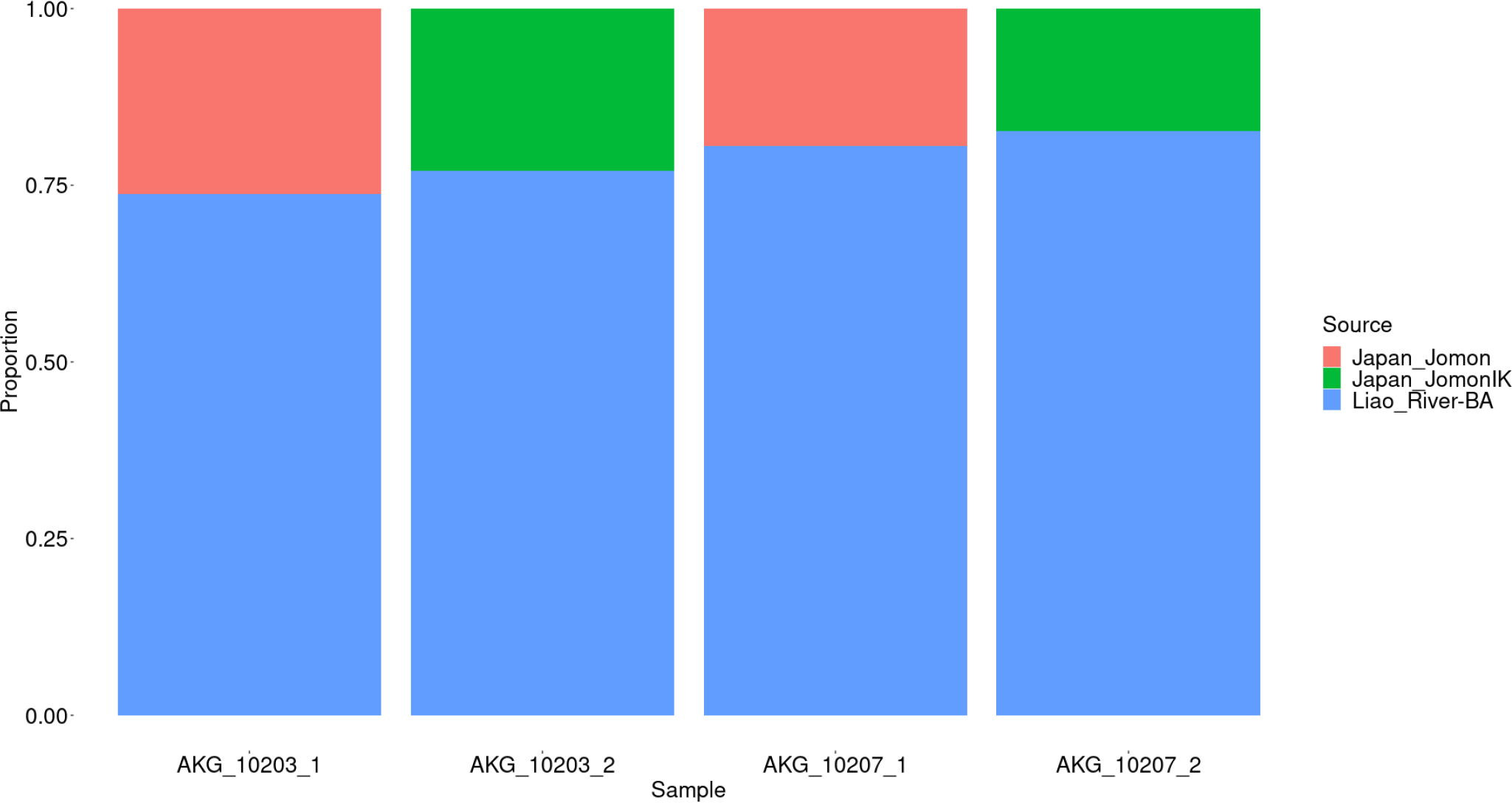
qpAdm results of all the working models of Korea-TK_2. The appendix numbers indicate the multiple models.

**Fig. S15:**
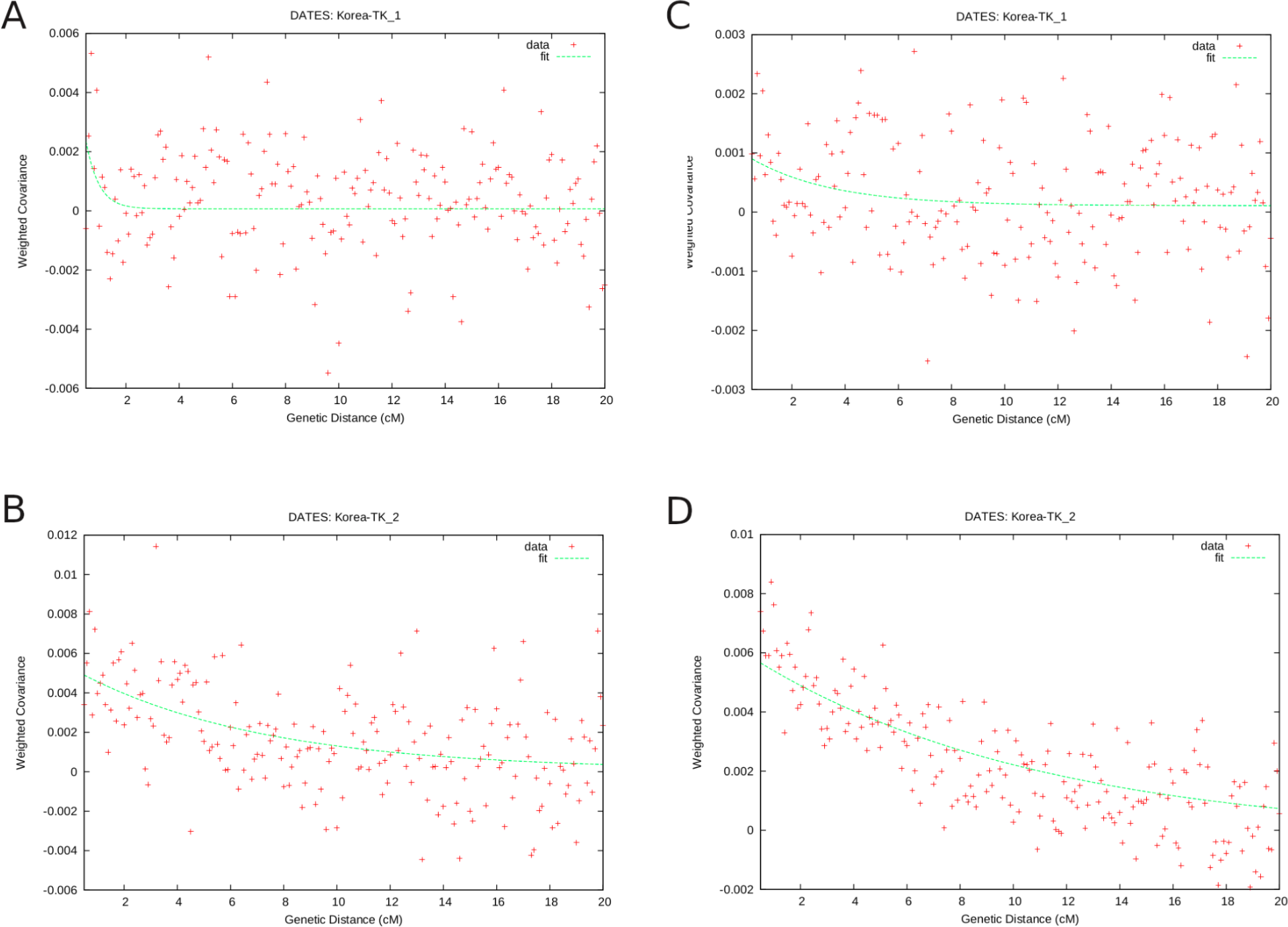
DATES analysis: A) Korea-TK_1 testing for Liao-River_BA and Japan_Jomon B) Korea-TK_2 testing for Liao-River_BA and Japan_Jomon C) Korea-TK_1 testing for Yellow-River_LBIA and Japan_Jomon D) Korea-TK_2 testing for Yellow-River_LBIA and Japan_Jomon.

**Fig. S16:**
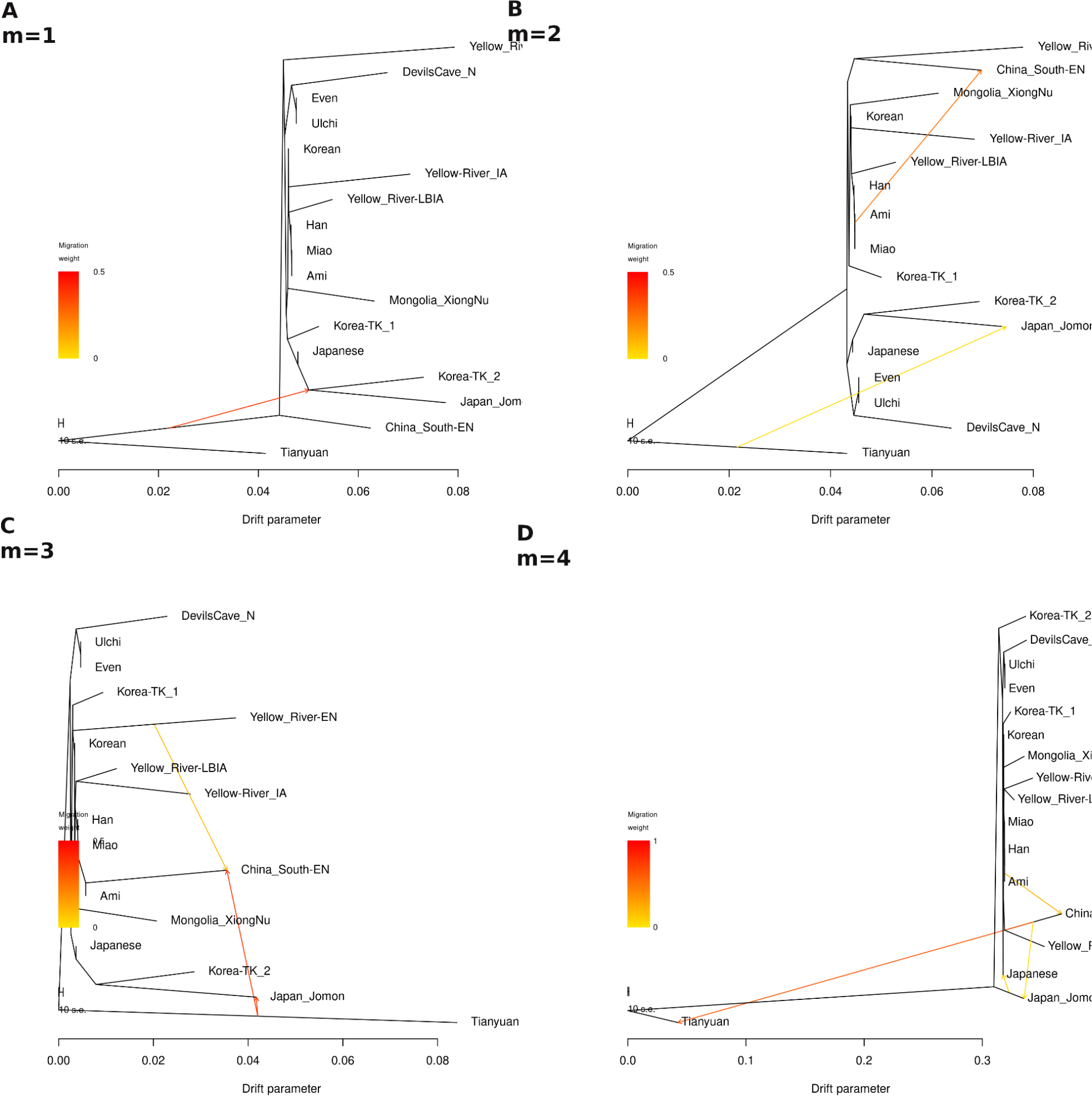
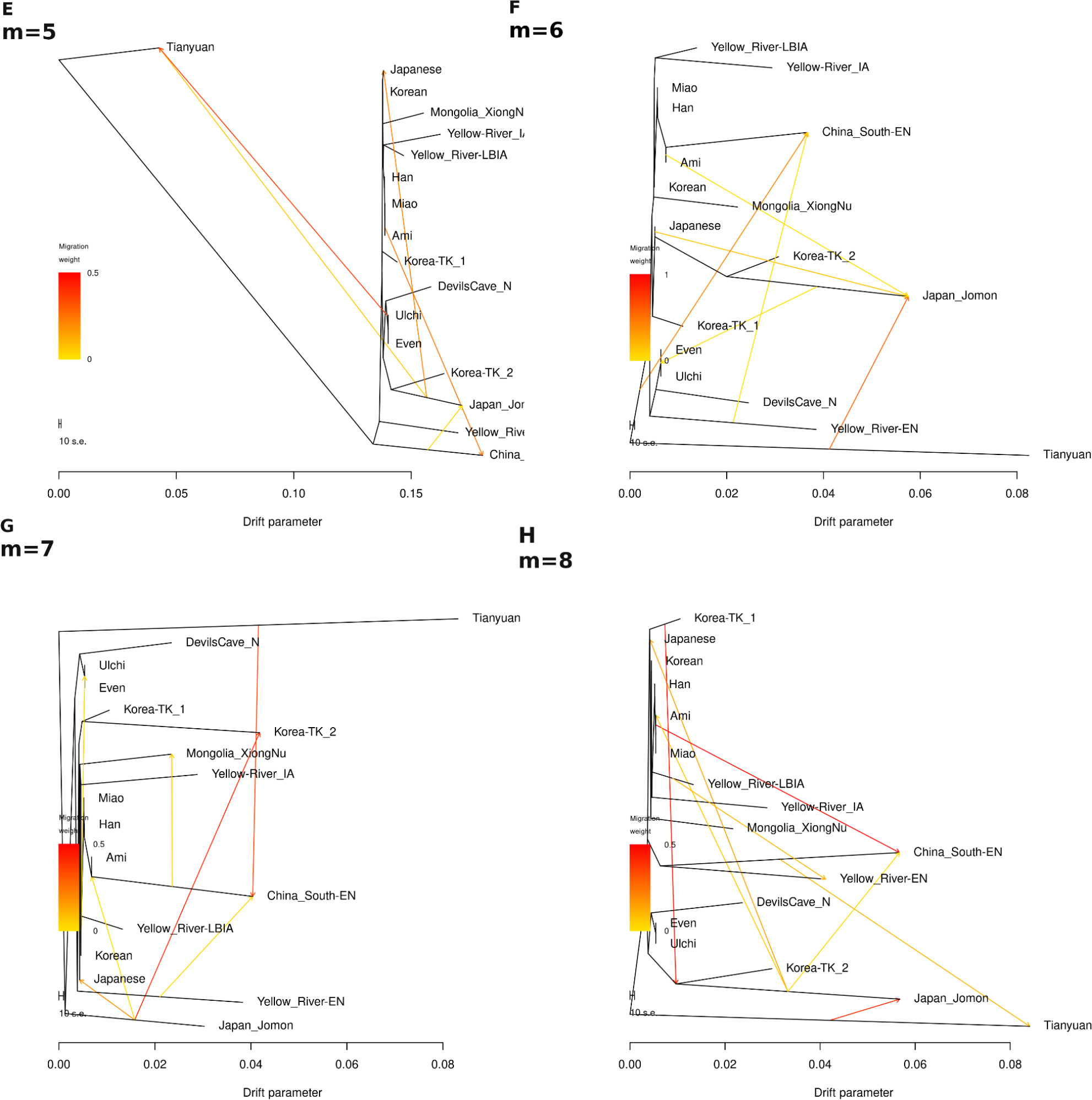
Treemix plots with the 15 North and East Asian populations and the two ancient Korean subgroups (Korea-TK_1, Korea-TK_2) modeled using one to eight migration events (m).

**Fig. S17:**
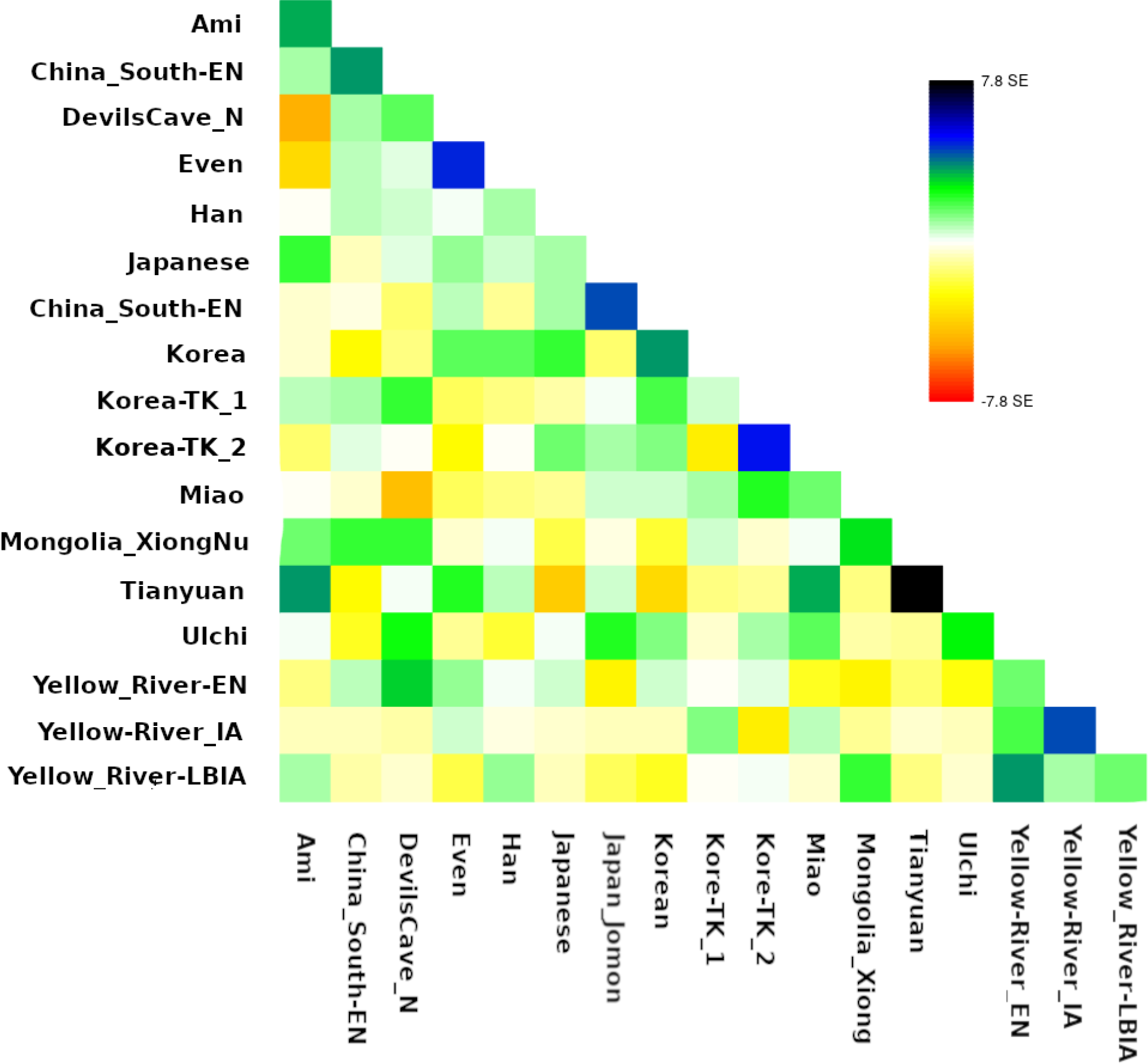
Treemix residue of the analysis using seven migrations

**Fig. S18:**
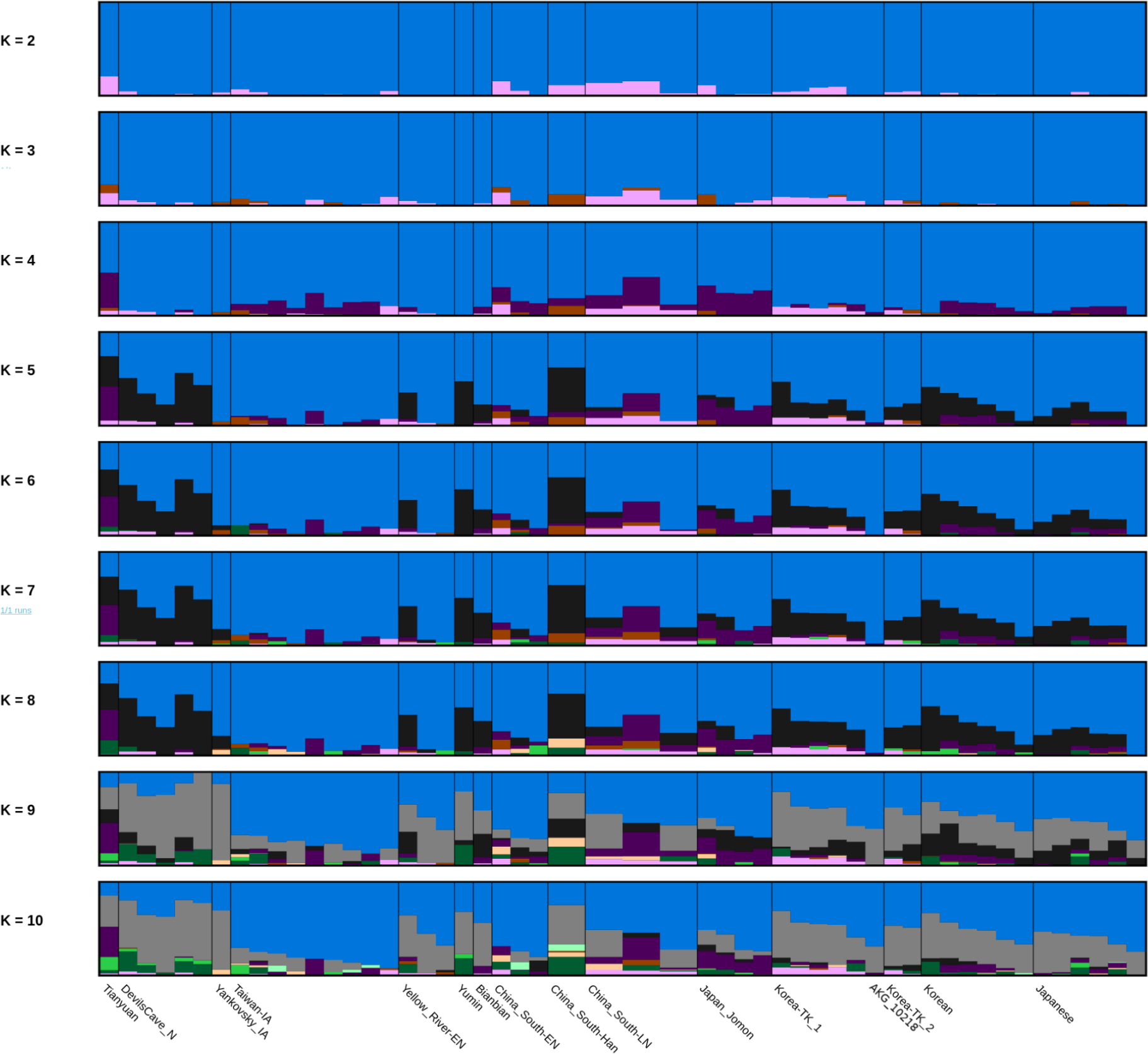
Admixture analysis of X chromosome SNPs with plots from K2 to K10.

**Figure S19:**
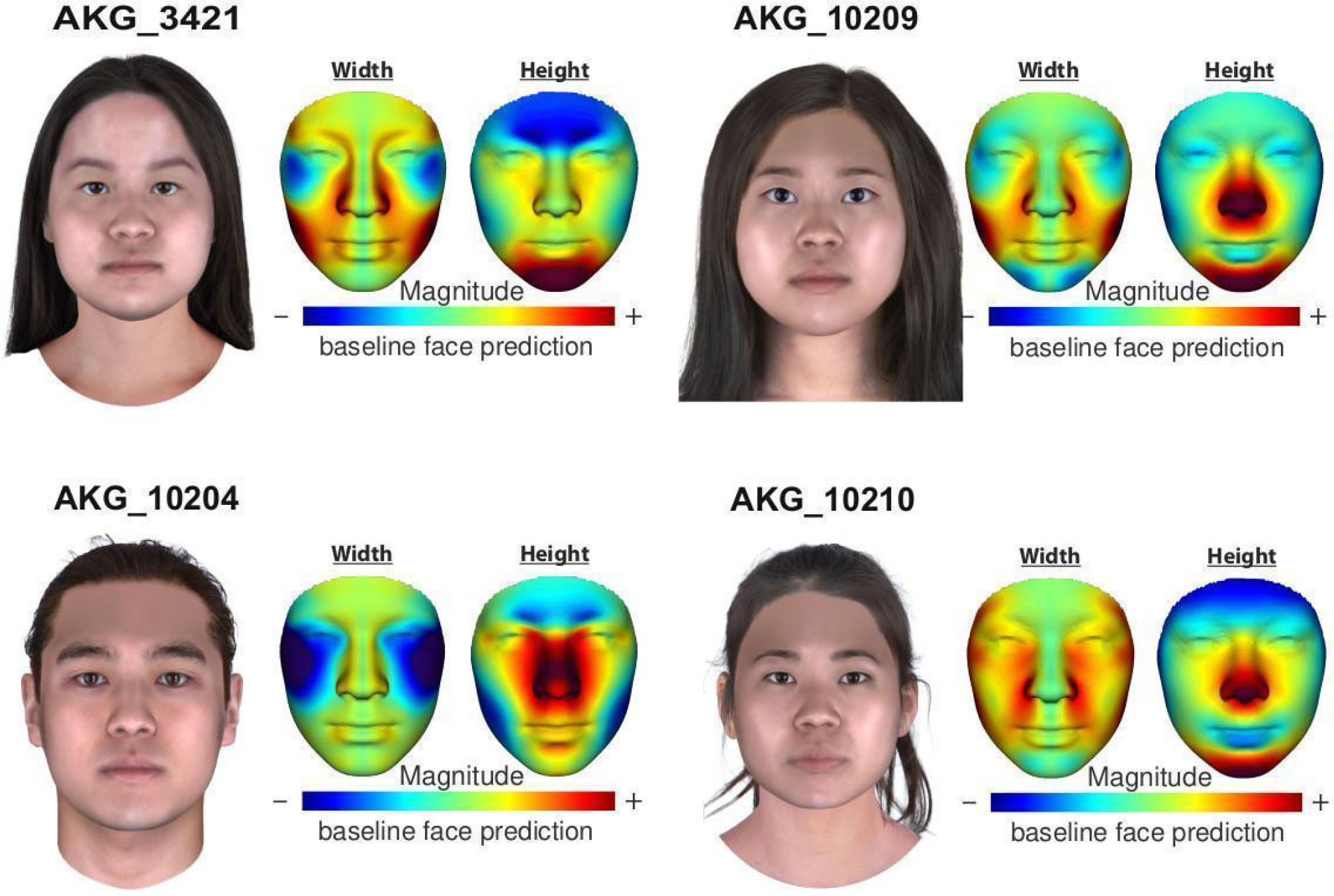
SNP-based facial predictions using a commercial program Snapshot® from Parabon NanoLabs (<u>h ttps://snapshot.parabon-nanolabs.com/</u>). Face morphology differences are emphasized relative to a baseline face prediction made using sex and ancestry. The facial reconstruction has been performed with the imputed calls dataset.

**Table S11:** Dates results: the dates are shown according to the admixtures suggested by the qpAdm results and only the ones showing credible curves.

| Admixed population | Pop A | Pop B | Generation to admixture | CI |
| --- | --- | --- | --- | --- |
| Korea-TK_1 | Jomon | YRBIA | 24.914 | 18.543 |
| Korea-TK_1 | Jomon | LRBA | 1097.120 | 416.527 |
| Korea-TK_2 | Jomon | YRBIA | 7.914 | 3.835 |
| Korea-TK_2 | Jomon | LRBA | 13.717 | 7.789 |

**Table S16:** Hirisplex prediction of the eye color. Blue SNPs have been predicted with the SNPs from the 1000 genomes database, while SNPs in black with the ones overlapping the Korean database with the 1000 genomes database.

| SNP | Allele | AKG_10<br>203 | AKG_102<br>04 | AKG_102<br>07 | AKG_102<br>09 | AKG_102<br>10 | AKG_102<br>18 | AKG_342<br>0 | AKG_342<br>1 |
| --- | --- | --- | --- | --- | --- | --- | --- | --- | --- |
| rs12913832 | T | 2 | 2 | 2 | 2 | 2 | 2 | 2 | 2 |
| rs1800407 | A | 0 | 0 | 0 | 0 | 0 | 0 | 0 | 0 |
| rs12896399 | T | 0 | 1 | 0 | 0 | 2 | 0 | 1 | 0 |
| rs16891982 | C | 2 | 2 | 2 | 2 | 2 | 2 | 2 | 2 |
| rs1393350 | T | 0 | 0 | 0 | 0 | 0 | 0 | 0 | 0 |
| rs12203592 | T | 0 | 0 | 0 | 0 | 1 | 0 | 0 | 0 |
| p-value |  | 0,998 | 0,997 | 0,998 | 0,998 | 0,993 | 0,998 | 0,997 | 0,998 |
| Result: |  | Brown<br>eyes | Brown<br>eyes | Brown<br>eyes | Brown<br>eyes | Brown<br>eyes | Brown<br>eyes | Brown<br>eyes | Brown<br>eyes |

**Table S17:** Hirisplex prediction of the hair color. Blue SNPs have been imputed with the SNPs from the 1000 genomes database, while SNPs in black with the ones overlapping the Korean database with the 1000 genomes database.

| SNP | Allele | AKG_10<br>203 | AKG_10<br>204 | AKG_10<br>207 | AKG_10<br>209 | AKG_10<br>210 | AKG_10<br>218 | AKG_34<br>20 | AKG_34<br>21 |
| --- | --- | --- | --- | --- | --- | --- | --- | --- | --- |
| rs312262906 | A | NA | NA | NA | NA | NA | NA | NA | NA |
| rs11547464 | A | 0 | 0 | 0 | 0 | 0 | 1 | 0 | 0 |
| rs885479 | T | 1 | 0 | 1 | 1 | 2 | 2 | 1 | 1 |
| rs1805008 | T | 0 | 0 | 0 | 0 | 0 | 0 | 0 | 0 |
| rs1805005 | T | 0 | 0 | 0 | 0 | 0 | 0 | 0 | 0 |
| rs1805006 | A | 0 | 0 | 0 | 0 | 0 | 0 | 0 | 0 |
| rs1805007 | T | 0 | 0 | 0 | 0 | 0 | 0 | 0 | 0 |
| rs1805009 | C | 0 | 0 | 0 | 0 | 0 | 0 | 0 | 0 |
| rs201326893 | A | NA | NA | NA | NA | NA | NA | NA | NA |
| rs2228479 | A | 0 | 1 | 0 | 0 | 0 | 0 | 0 | 0 |
| rs1110400 | C | 0 | 0 | 0 | 0 | 0 | 0 | 0 | 0 |
| rs28777 | C | 1 | 2 | 2 | 2 | 2 | 2 | 2 | 1 |
| rs16891982 | C | 2 | 2 | 2 | 2 | 2 | 2 | 2 | 2 |
| rs12821256 | G | 0 | 0 | 0 | 0 | 0 | 0 | 0 | 0 |
| rs4959270 | A | 0 | 0 | 1 | 0 | 0 | 0 | 0 | 1 |
| rs12203592 | T | 0 | 0 | 0 | 0 | 0 | 0 | 0 | 0 |
| rs1042602 | T | 0 | 0 | 0 | 0 | 0 | 0 | 0 | 0 |
| rs1800407 | A | 0 | 0 | 0 | 0 | 0 | 0 | 0 | 0 |
| rs2402130 | G | 1 | 0 | 1 | 1 | 0 | 1 | 0 | 0 |
| rs12913832 | T | 2 | 2 | 2 | 2 | 2 | 2 | 2 | 2 |
| rs2378249 | C | 0 | 0 | 0 | 0 | 1 | 0 | 0 | 2 |
| rs12896399 | T | 0 | 1 | 0 | 0 | 2 | 0 | 1 | 0 |
| rs1393350 | T | 0 | 0 | 0 | 0 | 0 | 0 | 0 | 0 |
| rs683 | G | 2 | 2 | 2 | 2 | 2 | 2 | 2 | 2 |
| rs3114908 | T | 1 | 1 | 1 | 0 | 1 | 1 | 1 | 0 |
| rs1800414 | C | 0 | 1 | 2 | 1 | 0 | 0 | 2 | 2 |
| rs10756819 | G | 1 | 1 | 2 | 2 | 1 | 0 | 1 | 2 |
| rs2238289 | C | 2 | 1 | 2 | 2 | 2 | 1 | 2 | 2 |
| rs17128291 | C | 1 | 0 | 0 | 0 | 0 | 0 | 1 | 0 |
| rs6497292 | C | 2 | 1 | 1 | 2 | 1 | 1 | 2 | 2 |
| rs1129038 | G | 2 | 2 | 2 | 1 | 2 | 2 | 2 | 2 |
| rs1667394 | C | 2 | 2 | 2 | 2 | 2 | 1 | 2 | 2 |
| rs1126809 | A | NA | NA | NA | NA | NA | NA | NA | NA |
| rs1470608 | A | NA | NA | NA | NA | NA | NA | NA | NA |
| rs1426654 | G | 2 | 2 | 2 | 2 | 2 | 2 | 2 | 2 |
| rs6119471 | C | NA | NA | NA | NA | NA | NA | NA | NA |
| rs1545397 | T | 2 | 2 | 2 | 2 | 2 | 2 | 2 | 2 |
| rs6059655 | T | 0 | 0 | 0 | 0 | 1 | 0 | 0 | 0 |
| rs12441727 | A | 0 | 0 | 0 | 0 | 1 | 0 | 0 | 0 |
| rs3212355 | A | 0 | 0 | 0 | 0 | 0 | 0 | 0 | 0 |
| rs8051733 | C | 0 | 1 | 0 | 0 | 0 | 0 | 0 | 0 |
| p-value |  | 0,997 | 0,997 | 0,998 | 0,998 | 0,999 | 0,777 | 0,998 | 0,996 |
| Prediction |  | Dark | Dark | Dark | Dark | Dark | Black | Dark | Dark |

**Table S18:** Hirisplex prediction of the skin color. Blue SNPs have been predicted with the SNPs from the 1000 genomes database, while SNPs in black with the ones overlapping the Korean database with the 1000 genomes database.

| SNP | Allele | AKG_102<br>03 | AKG_102<br>04 | AKG_102<br>07 | AKG_102<br>09 | AKG_102<br>10 | AKG_102<br>18 | AKG_342<br>0 | AKG_34<br>21 |
| --- | --- | --- | --- | --- | --- | --- | --- | --- | --- |
| rs312262906 | A | NA | NA | NA | NA | NA | NA | NA | NA |
| rs11547464 | A | 0 | 0 | 0 | 0 | 0 | 1 | 0 | 0 |
| rs885479 | T | 1 | 0 | 1 | 1 | 2 | 2 | 1 | 1 |
| rs1805008 | T | 0 | 0 | 0 | 0 | 0 | 0 | 0 | 0 |
| rs1805005 | T | 0 | 0 | 0 | 0 | 0 | 0 | 0 | 0 |
| rs1805006 | A | 0 | 0 | 0 | 0 | 0 | 0 | 0 | 0 |
| rs1805007 | T | 0 | 0 | 0 | 0 | 0 | 0 | 0 | 0 |
| rs1805009 | C | 0 | 0 | 0 | 0 | 0 | 0 | 0 | 0 |
| rs201326893 | A | NA | NA | NA | NA | NA | NA | NA | NA |
| rs2228479 | A | 0 | 1 | 0 | 0 | 0 | 0 | 0 | 0 |
| rs1110400 | C | 0 | 0 | 0 | 0 | 0 | 0 | 0 | 0 |
| rs28777 | C | 1 | 2 | 2 | 2 | 2 | 2 | 2 | 1 |
| rs16891982 | C | 2 | 2 | 2 | 2 | 2 | 2 | 2 | 2 |
| rs12821256 | G | 0 | 0 | 0 | 0 | 0 | 0 | 0 | 0 |
| rs4959270 | A | 0 | 0 | 1 | 0 | 0 | 0 | 0 | 1 |
| rs12203592 | T | 0 | 0 | 0 | 0 | 0 | 0 | 0 | 0 |
| rs1042602 | T | 0 | 0 | 0 | 0 | 0 | 0 | 0 | 0 |
| rs1800407 | A | 0 | 0 | 0 | 0 | 0 | 0 | 0 | 0 |
| rs2402130 | G | 1 | 0 | 1 | 1 | 0 | 1 | 0 | 0 |
| rs12913832 | T | 2 | 2 | 2 | 2 | 2 | 2 | 2 | 2 |
| rs2378249 | C | 0 | 0 | 0 | 0 | 1 | 0 | 0 | 2 |
| rs12896399 | T | 0 | 1 | 0 | 0 | 2 | 0 | 1 | 0 |
| rs1393350 | T | 0 | 0 | 0 | 0 | 0 | 0 | 0 | 0 |
| rs683 | G | 2 | 2 | 2 | 2 | 2 | 2 | 2 | 2 |
| rs3114908 | T | 1 | 1 | 1 | 0 | 1 | 1 | 1 | 0 |
| rs1800414 | C | 0 | 1 | 2 | 1 | 0 | 0 | 2 | 2 |
| rs10756819 | G | 1 | 1 | 2 | 2 | 1 | 0 | 1 | 2 |
| rs2238289 | C | 2 | 1 | 2 | 2 | 2 | 1 | 2 | 2 |
| rs17128291 | C | 1 | 0 | 0 | 0 | 0 | 0 | 1 | 0 |
| rs6497292 | C | 2 | 1 | 1 | 2 | 1 | 1 | 2 | 2 |
| rs1129038 | G | 2 | 2 | 2 | 1 | 2 | 2 | 2 | 2 |
| rs1667394 | C | 2 | 2 | 2 | 2 | 2 | 1 | 2 | 2 |
| rs1126809 | A | NA | NA | NA | NA | NA | NA | NA | NA |
| rs1470608 | A | NA | NA | NA | NA | NA | NA | NA | NA |
| rs1426654 | G | 2 | 2 | 2 | 2 | 2 | 2 | 2 | 2 |
| rs6119471 | C | NA | NA | NA | NA | NA | NA | NA | NA |
| rs1545397 | T | 2 | 2 | 2 | 2 | 2 | 2 | 2 | 2 |
| rs6059655 | T | 0 | 0 | 0 | 0 | 1 | 0 | 0 | 0 |
| rs12441727 | A | 0 | 0 | 0 | 0 | 1 | 0 | 0 | 0 |
| rs3212355 | A | 0 | 0 | 0 | 0 | 0 | 0 | 0 | 0 |
| rs8051733 | C | 0 | 1 | 0 | 0 | 0 | 0 | 0 | 0 |
| p-value |  | 0,753 | 0,955 | 0,837 | 0,685 | 0,818 | 0,916 | 0,913 | 0,994 |
| Prediction |  | Dark | Dark | Intermed<br>iate | Dark | Dark | Dark | Interme<br>diate | Interme<br>diate |

## References

1. C.-C. Wang, H.-Y. Yeh, A. N. Popov, H.-Q. Zhang, H. Matsumura, K. Sirak, O. Cheronet, A. Kovalev, N. Rohland, A. M. Kim, S. Mallick, R. Bernardos, D. Tumen, J. Zhao, Y.-C. Liu, J.-Y. Liu, M. Mah, K. Wang, Z. Zhang, N. Adamski, N. Broomandkhoshbacht, K. Callan, F. Candilio, K. S. D. Carlson, B. J. Culleton, L. Eccles, S. Freilich, D. Keating, A. M. Lawson, K. Mandl, M. Michel, J. Oppenheimer, K. T. Özdoğan, K. Stewardson, S. Wen, S. Yan, F. Zalzala, R. Chuang, C.-J. Huang, H. Looh, C.-C. Shiung, Y. G. Nikitin, A. V. Tabarev, A. A. Tishkin, S. Lin, Z.-Y. Sun, X.-M. Wu, T.-L. Yang, X. Hu, L. Chen, H. Du, J. Bayarsaikhan, E. Mijiddorj, D. Erdenebaatar, T.-O. Iderkhangai, E. Myagmar, H. Kanzawa-Kiriyama, M. Nishino, K.-I. Shinoda, O. A. Shubina, J. Guo, W. Cai, Q. Deng, L. Kang, D. Li, D. Li, R. Lin, Nini, R. Shrestha, L.-X. Wang, L. Wei, G. Xie, H. Yao, M. Zhang, G. He, X. Yang, R. Hu, M. Robbeets, S. Schiffels, D. J. Kennett, L. Jin, H. Li, J. Krause, R. Pinhasi, D. Reich, Genomic Insights into the Formation of Human Populations in East Asia. Nature (2021), doi:10.1038/s41586-021-03336-2.

2. C. Ning, T. Li, K. Wang, F. Zhang, T. Li, X. Wu, S. Gao, Q. Zhang, H. Zhang, M. J. Hudson, G. Dong, S. Wu, Y. Fang, C. Liu, C. Feng, W. Li, T. Han, R. Li, J. Wei, Y. Zhu, Y. Zhou, C.-C. Wang, S. Fan, Z. Xiong, Z. Sun, M. Ye, L. Sun, X. Wu, F. Liang, Y. Cao, X. Wei, H. Zhu, H. Zhou, J. Krause, M. Robbeets, C. Jeong, Y. Cui, Ancient genomes from northern China suggest links between subsistence changes and human migration. Nat. Commun. 11, 2700 (2020).

3. M. A. Yang, X. Fan, B. Sun, C. Chen, J. Lang, Y.-C. Ko, C.-H. Tsang, H. Chiu, T. Wang, Q. Bao, X. Wu, M. Hajdinjak, A. M.-S. Ko, M. Ding, P. Cao, R. Yang, F. Liu, B. Nickel, Q. Dai, X. Feng, L. Zhang, C. Sun, C. Ning, W. Zeng, Y. Zhao, M. Zhang, X. Gao, Y. Cui, D. Reich, M. Stoneking, Q. Fu, Ancient DNA indicates human population shifts and admixture in northern and southern China. Science (2020), doi:10.1126/science.aba0909.

4. H. McColl, F. Racimo, L. Vinner, F. Demeter, T. Gakuhari, J. V. Moreno-Mayar, G. van Driem, U. Gram Wilken, A. Seguin-Orlando, C. de la Fuente Castro, S. Wasef, R. Shoocongdej, V. Souksavatdy, T. Sayavongkhamdy, M. M. Saidin, M. E. Allentoft, T. Sato, A.-S. Malaspinas, F. A. Aghakhanian, T. Korneliussen, A. Prohaska, A. Margaryan, P. de Barros Damgaard, S. Kaewsutthi, P. Lertrit, T. M. H. Nguyen, H.-C. Hung, T. Minh Tran, H. Nghia Truong, G. H. Nguyen, S. Shahidan, K. Wiradnyana, H. Matsumae, N. Shigehara, M. Yoneda, H. Ishida, T. Masuyama, Y. Yamada, A. Tajima, H. Shibata, A. Toyoda, T. Hanihara, S. Nakagome, T. Deviese, A.-M. Bacon, P. Duringer, J.-L. Ponche, L. Shackelford, E. Patole-Edoumba, A. T. Nguyen, B. Bellina-Pryce, J.-C. Galipaud, R. Kinaston, H. Buckley, C. Pottier, S. Rasmussen, T. Higham, R. A. Foley, M. M. Lahr, L. Orlando, M. Sikora, M. E. Phipps, H. Oota, C. Higham, D. M. Lambert, E. Willerslev, The prehistoric peopling of Southeast Asia. Science. 361, 88–92 (2018).

5. T. Gakuhari, S. Nakagome, S. Rasmussen, M. E. Allentoft, T. Sato, T. Korneliussen, B. N. Chuinneagáin, H. Matsumae, K. Koganebuchi, R. Schmidt, S. Mizushima, O. Kondo, N. Shigehara, M. Yoneda, R. Kimura, H. Ishida, T. Masuyama, Y. Yamada, A. Tajima, H. Shibata, A. Toyoda, T. Tsurumoto, T. Wakebe, H. Shitara, T. Hanihara, E. Willerslev, M. Sikora, H. Oota, Ancient Jomon genome sequence analysis sheds light on migration patterns of early East Asian populations. Commun Biol. 3, 437 (2020).

6. C. Jeong, K. Wang, S. Wilkin, W. T. T. Taylor, B. K. Miller, J. H. Bemmann, R. Stahl, C. Chiovelli, F. Knolle, S. Ulziibayar, D. Khatanbaatar, D. Erdenebaatar, U. Erdenebat, A. Ochir, G. Ankhsanaa, C. Vanchigdash, B. Ochir, C. Munkhbayar, D. Tumen, A. Kovalev, N. Kradin, B. A. Bazarov, D. A. Miyagashev, P. B. Konovalov, E. Zhambaltarova, A. V. Miller, W. Haak, S. Schiffels, J. Krause, N. Boivin, M. Erdene, J. Hendy, C. Warinner, A Dynamic 6,000-Year Genetic History of Eurasia’s Eastern Steppe. Cell. 183, 890–904.e29 (2020).

7. P. de Barros Damgaard, R. Martiniano, J. Kamm, J. V. Moreno-Mayar, G. Kroonen, M. Peyrot, G. Barjamovic, S. Rasmussen, C. Zacho, N. Baimukhanov, V. Zaibert, V. Merz, A. Biddanda, I. Merz, V. Loman, V. Evdokimov, E. Usmanova, B. Hemphill, A. Seguin-Orlando, F. E. Yediay, I. Ullah, K.-G. Sjögren, K. H. Iversen, J. Choin, C. de la Fuente, M. Ilardo, H. Schroeder, V. Moiseyev, A. Gromov, A. Polyakov, S. Omura, S. Y. Senyurt, H. Ahmad, C. McKenzie, A. Margaryan, A. Hameed, A. Samad, N. Gul, M. H. Khokhar, O. I. Goriunova, V. I. Bazaliiskii, J. Novembre, A. W. Weber, L. Orlando, M. E. Allentoft, R. Nielsen, K. Kristiansen, M. Sikora, A. K. Outram, R. Durbin, E. Willerslev, The first horse herders and the impact of early Bronze Age steppe expansions into Asia. Science. 360 (2018), doi:10.1126/science.aar7711.

8. X. Mao, H. Zhang, S. Qiao, Y. Liu, F. Chang, P. Xie, M. Zhang, T. Wang, M. Li, P. Cao, R. Yang, F. Liu, Q. Dai, X. Feng, W. Ping, C. Lei, J. W. Olsen, E. A. Bennett, Q. Fu, The deep population history of northern East Asia from the Late Pleistocene to the Holocene. Cell (2021), doi:10.1016/j.cell.2021.04.040.

9. J. Kim, S. Jeon, J.-P. Choi, A. Blazyte, Y. Jeon, J.-I. Kim, J. Ohashi, K. Tokunaga, S. Sugano, S. Fucharoen, F. Al-Mulla, J. Bhak, The Origin and Composition of Korean Ethnicity Analyzed by Ancient and Present-Day Genome Sequences. Genome Biol. Evol. 12, 553–565 (2020).

10. S. Jeon, Y. Bhak, Y. Choi, Y. Jeon, S. Kim, J. Jang, J. Jang, A. Blazyte, C. Kim, Y. Kim, J. Shim, N. Kim, Y. J. Kim, S. G. Park, J. Kim, Y. S. Cho, Y. Park, H.-M. Kim, B.-C. Kim, N.-H. Park, E.-S. Shin, B. C. Kim, D. Bolser, A. Manica, J. S. Edwards, G. Church, S. Lee, J. Bhak, Korean Genome Project: 1094 Korean personal genomes with clinical information. Science Advances. 6, eaaz7835 (2020).

11. H.-J. Jin, C. Tyler-Smith, W. Kim, The peopling of Korea revealed by analyses of mitochondrial DNA and Y-chromosomal markers. PLoS One. 4, e4210 (2009).

12. R. M. Gutaker, S. C. Groen, E. S. Bellis, J. Y. Choi, I. S. Pires, R. K. Bocinsky, E. R. Slayton, O. Wilkins, C. C. Castillo, S. Negrão, M. M. Oliveira, D. Q. Fuller, J. A. D. Guedes, J. R. Lasky, M. D. Purugganan, Genomic history and ecology of the geographic spread of rice. Nature Plants. 6, 492–502 (2020).

13. G. W. Crawford, G.-A. Lee, Agricultural origins in the Korean Peninsula. Antiquity. 77, 87–95 (2003).

14. S.-M. Ahn, The emergence of rice agriculture in Korea: archaeobotanical perspectives. Archaeol. Anthropol. Sci. 2, 89–98 (2010).

15. P. Golas, Donald B. Wagner: Iron and Steel in Ancient China. Bull. Indiana State Dep. Health. 82, 426–428 (1995).

16. W. Rostoker, B. Bronson, Pre-industrial iron: its technology and ethnology (bcin.ca, 1990).

17. S. Shoda, A comment on the Yayoi period dating controversy. Bulletin of the society for East Asian archaeology. 1, 1–7 (2007).

18. M. Peterson, P. Margulies, A brief history of Korea (2019).

19. G. L. Barnes, Ed., in State Formation in Korea: Historical and Archaeological Perspectives (Curzon/Routledg, London, 2001).

20. M. L. Antonio, Z. Gao, H. M. Moots, M. Lucci, F. Candilio, S. Sawyer, V. Oberreiter, D. Calderon, K. Devitofranceschi, R. C. Aikens, S. Aneli, F. Bartoli, A. Bedini, O. Cheronet, D. J. Cotter, D. M. Fernandes, G. Gasperetti, R. Grifoni, A. Guidi, F. La Pastina, E. Loreti, D. Manacorda, G. Matullo, S. Morretta, A. Nava, V. Fiocchi Nicolai, F. Nomi, C. Pavolini, M. Pentiricci, P. Pergola, M. Piranomonte, R. Schmidt, G. Spinola, A. Sperduti, M. Rubini, L. Bondioli, A. Coppa, R. Pinhasi, J. K. Pritchard, Ancient Rome: A genetic crossroads of Europe and the Mediterranean. Science. 366, 708–714 (2019).

21. K.-C. Shin, Relations between Kaya and Wa in the third to fourth centuries AD. Journal of East Asian Archaeology. 2, 112–122 (2000).

22. S. Sawyer, J. Krause, K. Guschanski, V. Savolainen, S. Pääbo, Temporal patterns of nucleotide misincorporations and DNA fragmentation in ancient DNA. PLoS One. 7, e34131 (2012).

23. M. Lipatov, K. Sanjeev, R. Patro, K. R. Veeramah, Maximum Likelihood Estimation of Biological Relatedness from Low Coverage Sequencing Data. bioRxiv (2015), p. 023374.

24. H. Ringbauer, J. Novembre, M. Steinrücken, Detecting runs of homozygosity from low-coverage ancient DNA. bioRxiv (2020), p. 2020.05.31.126912.

25. W. Haak, I. Lazaridis, N. Patterson, N. Rohland, S. Mallick, B. Llamas, G. Brandt, S. Nordenfelt, E. Harney, K. Stewardson, Q. Fu, A. Mittnik, E. Bánffy, C. Economou, M. Francken, S. Friederich, R. G. Pena, F. Hallgren, V. Khartanovich, A. Khokhlov, M. Kunst, P. Kuznetsov, H. Meller, O. Mochalov, V. Moiseyev, N. Nicklisch, S. L. Pichler, R. Risch, M. A. Rojo Guerra, C. Roth, A. Szécsényi-Nagy, J. Wahl, M. Meyer, J. Krause, D. Brown, D. Anthony, A. Cooper, K. W. Alt, D. Reich, Massive migration from the steppe was a source for Indo-European languages in Europe. Nature. 522, 207–211 (2015).

26. R. Martiniano, L. M. Cassidy, R. Ó’Maoldúin, R. McLaughlin, N. M. Silva, L. Manco, D. Fidalgo, T. Pereira, M. J. Coelho, M. Serra, J. Burger, R. Parreira, E. Moran, A. C. Valera, E. Porfirio, R. Boaventura, A. M. Silva, D. G. Bradley, The population genomics of archaeological transition in west Iberia: Investigation of ancient substructure using imputation and haplotype-based methods. PLoS Genet. 13, e1006852 (2017).

27. G. Renaud, V. Slon, A. T. Duggan, J. Kelso, Schmutzi: estimation of contamination and endogenous mitochondrial consensus calling for ancient DNA. Genome Biol. 16, 224 (2015).

28. H. Weissensteiner, D. Pacher, A. Kloss-Brandstätter, L. Forer, G. Specht, H.-J. Bandelt, F. Kronenberg, A. Salas, S. Schönherr, HaploGrep 2: mitochondrial haplogroup classification in the era of high-throughput sequencing. Nucleic Acids Res. 44, W58–63 (2016).

29. F. Mizuno, J. Gojobori, M. Kumagai, H. Baba, Y. Taniguchi, O. Kondo, M. Matsushita, T. Matsushita, F. Matsuda, K. Higasa, M. Hayashi, L. Wang, K. Kurosaki, S. Ueda, Population dynamics in the Japanese Archipelago since the Pleistocene revealed by the complete mitochondrial genome sequences. Sci. Rep. 11, 12018 (2021).

30. S. S. Choi, K. H. Park, D. E. Nam, T. H. Kang, K. W. Chung, Y-chromosome haplogrouping for Asians using Y-SNP target sequencing. Forensic Science International: Genetics Supplement Series. 6, e235–e237 (2017).

31. I. Lazaridis, N. Patterson, A. Mittnik, G. Renaud, S. Mallick, K. Kirsanow, P. H. Sudmant, J. G. Schraiber, S. Castellano, M. Lipson, B. Berger, C. Economou, R. Bollongino, Q. Fu, K. I. Bos, S. Nordenfelt, H. Li, C. de Filippo, K. Prüfer, S. Sawyer, C. Posth, W. Haak, F. Hallgren, E. Fornander, N. Rohland, D. Delsate, M. Francken, J.-M. Guinet, J. Wahl, G. Ayodo, H. A. Babiker, G. Bailliet, E. Balanovska, O. Balanovsky, R. Barrantes, G. Bedoya, H. Ben-Ami, J. Bene, F. Berrada, C. M. Bravi, F. Brisighelli, G. B. J. Busby, F. Cali, M. Churnosov, D. E. C. Cole, D. Corach, L. Damba, G. van Driem, S. Dryomov, J.-M. Dugoujon, S. A. Fedorova, I. Gallego Romero, M. Gubina, M. Hammer, B. M. Henn, T. Hervig, U. Hodoglugil, A. R. Jha, S. Karachanak-Yankova, R. Khusainova, E. Khusnutdinova, R. Kittles, T. Kivisild, W. Klitz, V. Kučinskas, A. Kushniarevich, L. Laredj, S. Litvinov, T. Loukidis, R. W. Mahley, B. Melegh, E. Metspalu, J. Molina, J. Mountain, K. Näkkäläjärvi, D. Nesheva, T. Nyambo, L. Osipova, J. Parik, F. Platonov, O. Posukh, V. Romano, F. Rothhammer, I. Rudan, R. Ruizbakiev, H. Sahakyan, A. Sajantila, A. Salas, E. B. Starikovskaya, A. Tarekegn, D. Toncheva, S. Turdikulova, I. Uktveryte, O. Utevska, R. Vasquez, M. Villena, M. Voevoda, C. A. Winkler, L. Yepiskoposyan, P. Zalloua, T. Zemunik, A. Cooper, C. Capelli, M. G. Thomas, A. Ruiz-Linares, S. A. Tishkoff, L. Singh, K. Thangaraj, R. Villems, D. Comas, R. Sukernik, M. Metspalu, M. Meyer, E. E. Eichler, J. Burger, M. Slatkin, S. Pääbo, J. Kelso, D. Reich, J. Krause, Ancient human genomes suggest three ancestral populations for present-day Europeans. Nature. 513, 409–413 (2014).

32. V. M. Narasimhan, N. Patterson, P. Moorjani, N. Rohland, R. Bernardos, S. Mallick, I. Lazaridis, N. Nakatsuka, I. Olalde, M. Lipson, A. M. Kim, L. M. Olivieri, A. Coppa, M. Vidale, J. Mallory, V. Moiseyev, E. Kitov, J. Monge, N. Adamski, N. Alex, N. Broomandkhoshbacht, F. Candilio, K. Callan, O. Cheronet, B. J. Culleton, M. Ferry, D. Fernandes, S. Freilich, B. Gamarra, D. Gaudio, M. Hajdinjak, É. Harney, T. K. Harper, D. Keating, A. M. Lawson, M. Mah, K. Mandl, M. Michel, M. Novak, J. Oppenheimer, N. Rai, K. Sirak, V. Slon, K. Stewardson, F. Zalzala, Z. Zhang, G. Akhatov, A. N. Bagashev, A. Bagnera, B. Baitanayev, J. Bendezu-Sarmiento, A. A. Bissembaev, G. L. Bonora, T. T. Chargynov, T. Chikisheva, P. K. Dashkovskiy, A. Derevianko, M. Dobeš, K. Douka, N. Dubova, M. N. Duisengali, D. Enshin, A. Epimakhov, A. V. Fribus, D. Fuller, A. Goryachev, A. Gromov, S. P. Grushin, B. Hanks, M. Judd, E. Kazizov, A. Khokhlov, A. P. Krygin, E. Kupriyanova, P. Kuznetsov, D. Luiselli, F. Maksudov, A. M. Mamedov, T. B. Mamirov, C. Meiklejohn, D. C. Merrett, R. Micheli, O. Mochalov, S. Mustafokulov, A. Nayak, D. Pettener, R. Potts, D. Razhev, M. Rykun, S. Sarno, T. M. Savenkova, K. Sikhymbaeva, S. M. Slepchenko, O. A. Soltobaev, N. Stepanova, S. Svyatko, K. Tabaldiev, M. Teschler-Nicola, A. A. Tishkin, V. V. Tkachev, S. Vasilyev, P. Velemínský, D. Voyakin, A. Yermolayeva, M. Zahir, V. S. Zubkov, A. Zubova, V. S. Shinde, C. Lalueza-Fox, M. Meyer, D. Anthony, N. Boivin, K. Thangaraj, D. J. Kennett, M. Frachetti, R. Pinhasi, D. Reich, The formation of human populations in South and Central Asia. Science. 365 (2019), doi:10.1126/science.aat7487.

33. K.-I. Yoshiura, A. Kinoshita, T. Ishida, A. Ninokata, T. Ishikawa, T. Kaname, M. Bannai, K. Tokunaga, S. Sonoda, R. Komaki, M. Ihara, V. A. Saenko, G. K. Alipov, I. Sekine, K. Komatsu, H. Takahashi, M. Nakashima, N. Sosonkina, C. K. Mapendano, M. Ghadami, M. Nomura, D.-S. Liang, N. Miwa, D.-K. Kim, A. Garidkhuu, N. Natsume, T. Ohta, H. Tomita, A. Kaneko, M. Kikuchi, G. Russomando, K. Hirayama, M. Ishibashi, A. Takahashi, N. Saitou, J. C. Murray, S. Saito, Y. Nakamura, N. Niikawa, A SNP in the ABCC11 gene is the determinant of human earwax type. Nat. Genet. 38, 324–330 (2006).

34. N. S. Enattah, T. Sahi, E. Savilahti, J. D. Terwilliger, L. Peltonen, I. Järvelä, Identification of a variant associated with adult-type hypolactasia. Nat. Genet. 30, 233–237 (2002).

35. C. Endo, T. A. Johnson, R. Morino, K. Nakazono, S. Kamitsuji, M. Akita, M. Kawajiri, T. Yamasaki, A. Kami, Y. Hoshi, A. Tada, K. Ishikawa, M. Hine, M. Kobayashi, N. Kurume, Y. Tsunemi, N. Kamatani, M. Kawashima, Genome-wide association study in Japanese females identifies fifteen novel skin-related trait associations. Sci. Rep. 8, 8974 (2018).

36. A. Fujimoto, J. Ohashi, N. Nishida, T. Miyagawa, Y. Morishita, T. Tsunoda, R. Kimura, K. Tokunaga, A replication study confirmed the EDAR gene to be a major contributor to population differentiation regarding head hair thickness in Asia. Hum. Genet. 124, 179–185 (2008).

37. L. Chaitanya, K. Breslin, S. Zuñiga, L. Wirken, E. Pośpiech, M. Kukla-Bartoszek, T. Sijen, P. de Knijff, F. Liu, W. Branicki, M. Kayser, S. Walsh, The HIrisPlex-S system for eye, hair and skin colour prediction from DNA: Introduction and forensic developmental validation. Forensic Sci. Int. Genet. 35, 123–135 (2018).

38. S.-K. Jung, J. H. Lee, H. Kakizaki, D. Jee, Prevalence of myopia and its association with body stature and educational level in 19-year-old male conscripts in seoul, South Korea. Invest. Ophthalmol. Vis. Sci. 53, 5579–5583 (2012).

39. C. Impraim, G. Wang, A. Yoshida, Structural mutation in a major human aldehyde dehydrogenase gene results in loss of enzyme activity. Am. J. Hum. Genet. 34, 837–841 (1982).

40. H. Jörnvall, J. O. Höög, Nomenclature of alcohol dehydrogenases. Alcohol Alcohol. 30, 153–161 (1995).

41. R. Li, F. F. Brockschmidt, A. K. Kiefer, H. Stefansson, D. R. Nyholt, K. Song, S. H. Vermeulen, S. Kanoni, D. Glass, S. E. Medland, M. Dimitriou, D. Waterworth, J. Y. Tung, F. Geller, S. Heilmann, A. M. Hillmer, V. Bataille, S. Eigelshoven, S. Hanneken, S. Moebus, C. Herold, M. den Heijer, G. W. Montgomery, P. Deloukas, N. Eriksson, A. C. Heath, T. Becker, P. Sulem, M. Mangino, P. Vollenweider, T. D. Spector, G. Dedoussis, N. G. Martin, L. A. Kiemeney, V. Mooser, K. Stefansson, D. A. Hinds, M. M. Nöthen, J. B. Richards, Six novel susceptibility Loci for early-onset androgenetic alopecia and their unexpected association with common diseases. PLoS Genet. 8, e1002746 (2012).

42. M. Larena, F. Sanchez-Quinto, P. Sjödin, J. McKenna, C. Ebeo, R. Reyes, O. Casel, J.-Y. Huang, K. P. Hagada, D. Guilay, J. Reyes, F. P. Allian, V. Mori, L. S. Azarcon, A. Manera, C. Terando, L. Jamero Jr, G. Sireg, R. Manginsay-Tremedal, M. S. Labos, R. D. Vilar, A. Latiph, R. L. Saway, E. Marte, P. Magbanua, A. Morales, I. Java, R. Reveche, B. Barrios, E. Burton, J. C. Salon, M. J. T. Kels, A. Albano, R. B. Cruz-Angeles, E. Molanida, L. Granehäll, M. Vicente, H. Edlund, J.-H. Loo, J. Trejaut, S. Y. W. Ho, L. Reid, H. Malmström, C. Schlebusch, K. Lambeck, P. Endicott, M. Jakobsson, Multiple migrations to the Philippines during the last 50,000 years. Proc. Natl. Acad. Sci. U. S. A. 118 (2021), doi:10.1073/pnas.2026132118.

43. K. Imamura, Prehistoric japan: New perspectives on insular east Asia (Routledge, London, England, 2016; https://www.taylorfrancis.com/books/9780203973424).

44. I. Nonaka, K. Minaguchi, N. Takezaki, Y-chromosomal binary haplogroups in the Japanese population and their relationship to 16 Y-STR polymorphisms. Ann. Hum. Genet. 71, 480–495 (2007).

45. N. Egami, Light on Japanese cultural origins from historical archaeology and legend. Japanese culture: its development and characteristics, 11–16 (1962).

46. W. Hong, Ancient Korea-japan relations: Dating the formative years of the Yamato kingdom (366-405 CE) by the samguk-sagi records and reinterpreting the related historical facts. Open area stud. j. 2, 12–29 (2009).

47. C.-C. Wang, H.-Y. Yeh, A. N. Popov, H.-Q. Zhang, H. Matsumura, K. Sirak, O. Cheronet, A. Kovalev, N. Rohland, A. M. Kim, R. Bernardos, D. Tumen, J. Zhao, Y.-C. Liu, J.-Y. Liu, M. Mah, S. Mallick, K. Wang, Z. Zhang, N. Adamski, N. Broomandkhoshbacht, K. Callan, B. J. Culleton, L. Eccles, A. M. Lawson, M. Michel, J. Oppenheimer, K. Stewardson, S. Wen, S. Yan, F. Zalzala, R. Chuang, C.-J. Huang, C.-C. Shiung, Y. G. Nikitin, A. V. Tabarev, A. A. Tishkin, S. Lin, Z.-Y. Sun, X.-M. Wu, T.-L. Yang, X. Hu, L. Chen, H. Du, J. Bayarsaikhan, E. Mijiddorj, D. Erdenebaatar, T.-O. Iderkhangai, E. Myagmar, H. Kanzawa-Kiriyama, M. Nishino, K.-I. Shinoda, O. A. Shubina, J. Guo, Q. Deng, L. Kang, D. Li, D. Li, R. Lin, W. Cai, R. Shrestha, L.-X. Wang, L. Wei, G. Xie, H. Yao, M. Zhang, G. He, X. Yang, R. Hu, M. Robbeets, S. Schiffels, D. J. Kennett, L. Jin, H. Li, J. Krause, R. Pinhasi, D. Reich, The Genomic Formation of Human Populations in East Asia. bioRxiv (2020), p. 2020.03.25.004606.

48. Q. Fu, M. Meyer, X. Gao, U. Stenzel, H. A. Burbano, J. Kelso, S. Pääbo, DNA analysis of an early modern human from Tianyuan Cave, China. Proc. Natl. Acad. Sci. U. S. A. 110, 2223–2227 (2013).

49. K. A. Sirak, D. M. Fernandes, O. Cheronet, M. Novak, B. Gamarra, T. Balassa, Z. Bernert, A. Cséki, J. Dani, J. Z. Gallina, G. Kocsis-Buruzs, I. Kővári, O. László, I. Pap, R. Patay, Z. Petkes, G. Szenthe, T. Szeniczey, T. Hajdu, R. Pinhasi, A minimally-invasive method for sampling human petrous bones from the cranial base for ancient DNA analysis. Biotechniques. 62, 283–289 (2017).

50. N. Rohland, I. Glocke, A. Aximu-Petri, M. Meyer, Extraction of highly degraded DNA from ancient bones, teeth and sediments for high-throughput sequencing. Nat. Protoc. 13, 2447–2461 (2018).

51. C. Gamba, E. R. Jones, M. D. Teasdale, R. L. McLaughlin, G. Gonzalez-Fortes, V. Mattiangeli, L. Domboróczki, I. Kővári, I. Pap, A. Anders, A. Whittle, J. Dani, P. Raczky, T. F. G. Higham, M. Hofreiter, D. G. Bradley, R. Pinhasi, Genome flux and stasis in a five millennium transect of European prehistory. Nat. Commun. 5, 5257 (2014).

52. M. Meyer, M. Kircher, Illumina sequencing library preparation for highly multiplexed target capture and sequencing. Cold Spring Harb. Protoc. 2010, db.prot5448 (2010).

53. M. Martin, Cutadapt removes adapter sequences from high-throughput sequencing reads. EMBnet.journal. 17, 10–12 (2011).

54. H. Li, R. Durbin, Fast and accurate short read alignment with Burrows-Wheeler transform. Bioinformatics. 25, 1754–1760 (2009).

55. Picard-tools, (available at http://broadinstitute.github.io/picard.).

56. K. Okonechnikov, A. Conesa, F. García-Alcalde, Qualimap 2: advanced multi-sample quality control for high-throughput sequencing data. Bioinformatics. 32, 292–294 (2016).

57. H. Jónsson, A. Ginolhac, M. Schubert, P. L. F. Johnson, L. Orlando, mapDamage2.0: fast approximate Bayesian estimates of ancient DNA damage parameters. Bioinformatics. 29, 1682–1684 (2013).

58. S. Schiffels, W. Haak, P. Paajanen, B. Llamas, E. Popescu, L. Loe, R. Clarke, A. Lyons, R. Mortimer, D. Sayer, C. Tyler-Smith, A. Cooper, R. Durbin, Iron Age and Anglo-Saxon genomes from East England reveal British migration history. Nat. Commun. 7, 10408 (2016).

59. M. Sikora, V. V. Pitulko, V. C. Sousa, M. E. Allentoft, L. Vinner, S. Rasmussen, A. Margaryan, P. de Barros Damgaard, C. de la Fuente, G. Renaud, M. A. Yang, Q. Fu, I. Dupanloup, K. Giampoudakis, D. Nogués-Bravo, C. Rahbek, G. Kroonen, M. Peyrot, H. McColl, S. V. Vasilyev, E. Veselovskaya, M. Gerasimova, E. Y. Pavlova, V. G. Chasnyk, P. A. Nikolskiy, A. V. Gromov, V. I. Khartanovich, V. Moiseyev, P. S. Grebenyuk, A. Y. Fedorchenko, A. I. Lebedintsev, S. B. Slobodin, B. A. Malyarchuk, R. Martiniano, M. Meldgaard, L. Arppe, J. U. Palo, T. Sundell, K. Mannermaa, M. Putkonen, V. Alexandersen, C. Primeau, N. Baimukhanov, R. S. Malhi, K.-G. Sjögren, K. Kristiansen, A. Wessman, A. Sajantila, M. M. Lahr, R. Durbin, R. Nielsen, D. J. Meltzer, L. Excoffier, E. Willerslev, The population history of northeastern Siberia since the Pleistocene. Nature. 570, 182–188 (2019).

60. M. Lipson, O. Cheronet, S. Mallick, N. Rohland, M. Oxenham, M. Pietrusewsky, T. O. Pryce, A. Willis, H. Matsumura, H. Buckley, K. Domett, G. H. Nguyen, H. H. Trinh, A. A. Kyaw, T. T. Win, B. Pradier, N. Broomandkhoshbacht, F. Candilio, P. Changmai, D. Fernandes, M. Ferry, B. Gamarra, E. Harney, J. Kampuansai, W. Kutanan, M. Michel, M. Novak, J. Oppenheimer, K. Sirak, K. Stewardson, Z. Zhang, P. Flegontov, R. Pinhasi, D. Reich, Ancient genomes document multiple waves of migration in Southeast Asian prehistory. Science. 361, 92–95 (2018).

61. V. Siska, E. R. Jones, S. Jeon, Y. Bhak, H.-M. Kim, Y. S. Cho, H. Kim, K. Lee, E. Veselovskaya, T. Balueva, M. Gallego-Llorente, M. Hofreiter, D. G. Bradley, A. Eriksson, R. Pinhasi, J. Bhak, A. Manica, Genome-wide data from two early Neolithic East Asian individuals dating to 7700 years ago. Sci Adv. 3, e1601877 (2017).

62. C. Jeong, A. T. Ozga, D. B. Witonsky, H. Malmström, H. Edlund, C. A. Hofman, R. W. Hagan, M. Jakobsson, C. M. Lewis, M. S. Aldenderfer, A. Di Rienzo, C. Warinner, Long-term genetic stability and a high-altitude East Asian origin for the peoples of the high valleys of the Himalayan arc. Proc. Natl. Acad. Sci. U. S. A. 113, 7485–7490 (2016).

63. H. Yu, M. A. Spyrou, M. Karapetian, S. Shnaider, R. Radzevičiūtė, K. Nägele, G. U. Neumann, S. Penske, J. Zech, M. Lucas, P. LeRoux, P. Roberts, G. Pavlenok, A. Buzhilova, C. Posth, C. Jeong, J. Krause, Paleolithic to Bronze Age Siberians Reveal Connections with First Americans and across Eurasia. Cell (2020), doi:10.1016/j.cell.2020.04.037.

64. I. Lazaridis, D. Nadel, G. Rollefson, D. C. Merrett, N. Rohland, S. Mallick, D. Fernandes, M. Novak, B. Gamarra, K. Sirak, S. Connell, K. Stewardson, E. Harney, Q. Fu, G. Gonzalez-Fortes, E. R. Jones, S. A. Roodenberg, G. Lengyel, F. Bocquentin, B. Gasparian, J. M. Monge, M. Gregg, V. Eshed, A.-S. Mizrahi, C. Meiklejohn, F. Gerritsen, L. Bejenaru, M. Blüher, A. Campbell, G. Cavalleri, D. Comas, P. Froguel, E. Gilbert, S. M. Kerr, P. Kovacs, J. Krause, D. McGettigan, M. Merrigan, D. A. Merriwether, S. O’Reilly, M. B. Richards, O. Semino, M. Shamoon-Pour, G. Stefanescu, M. Stumvoll, A. Tönjes, A. Torroni, J. F. Wilson, L. Yengo, N. A. Hovhannisyan, N. Patterson, R. Pinhasi, D. Reich, Genomic insights into the origin of farming in the ancient Near East. Nature. 536, 419–424 (2016).

65. T. S. Korneliussen, A. Albrechtsen, R. Nielsen, ANGSD: Analysis of Next Generation Sequencing Data. BMC Bioinformatics. 15, 356 (2014).

66. S. Andrews, FastQC: A Quality Control Tool for High Throughput Sequence Data (2010).

67. A. Ralf, D. Montiel González, K. Zhong, M. Kayser, Yleaf: Software for Human Y-Chromosomal Haplogroup Inference from Next-Generation Sequencing Data. Mol. Biol. Evol. 35, 1291–1294 (2018).

68. J. M. Monroy Kuhn, M. Jakobsson, T. Günther, Estimating genetic kin relationships in prehistoric populations. PLoS One. 13, e0195491 (2018).

69. N. Patterson, P. Moorjani, Y. Luo, S. Mallick, N. Rohland, Y. Zhan, T. Genschoreck, T. Webster, D. Reich, Ancient admixture in human history. Genetics. 192, 1065–1093 (2012).

70. R Core Team, R: A language and environment for statistical computing. R Foundation for Statistical Computing, Vienna, Austria (2013), (available at http://www.R-project.org/.).

71. D. H. Alexander, J. Novembre, K. Lange, Fast model-based estimation of ancestry in unrelated individuals. Genome Res. 19, 1655–1664 (2009).

72. S. Purcell, B. Neale, K. Todd-Brown, L. Thomas, M. A. R. Ferreira, D. Bender, J. Maller, P. Sklar, P. I. W. de Bakker, M. J. Daly, P. C. Sham, PLINK: a tool set for whole-genome association and population-based linkage analyses. Am. J. Hum. Genet. 81, 559–575 (2007).

73. A. A. Behr, K. Z. Liu, G. Liu-Fang, P. Nakka, S. Ramachandran, pong: fast analysis and visualization of latent clusters in population genetic data. Bioinformatics. 32, 2817–2823 (2016).

74. D. Reich, N. Patterson, D. Campbell, A. Tandon, S. Mazieres, N. Ray, M. V. Parra, W. Rojas, C. Duque, N. Mesa, L. F. García, O. Triana, S. Blair, A. Maestre, J. C. Dib, C. M. Bravi, G. Bailliet, D. Corach, T. Hünemeier, M. C. Bortolini, F. M. Salzano, M. L. Petzl-Erler, V. Acuña-Alonzo, C. Aguilar-Salinas, S. Canizales-Quinteros, T. Tusié-Luna, L. Riba, M. Rodríguez-Cruz, M. Lopez-Alarcón, R. Coral-Vazquez, T. Canto-Cetina, I. Silva-Zolezzi, J. C. Fernandez-Lopez, A. V. Contreras, G. Jimenez-Sanchez, M. J. Gómez-Vázquez, J. Molina, A. Carracedo, A. Salas, C. Gallo, G. Poletti, D. B. Witonsky, G. Alkorta-Aranburu, R. I. Sukernik, L. Osipova, S. A. Fedorova, R. Vasquez, M. Villena, C. Moreau, R. Barrantes, D. Pauls, L. Excoffier, G. Bedoya, F. Rothhammer, J.-M. Dugoujon, G. Larrouy, W. Klitz, D. Labuda, J. Kidd, K. Kidd, A. Di Rienzo, N. B. Freimer, A. L. Price, A. Ruiz-Linares, Reconstructing Native American population history. Nature. 488, 370–374 (2012).

75. J. K. Pickrell, J. K. Pritchard, Inference of population splits and mixtures from genome-wide allele frequency data. PLoS Genet. 8, e1002967 (2012).

76. P. Moorjani, S. Sankararaman, Q. Fu, M. Przeworski, N. Patterson, D. Reich, A genetic method for dating ancient genomes provides a direct estimate of human generation interval in the last 45,000 years. Proc. Natl. Acad. Sci. U. S. A. 113, 5652–5657 (2016).

77. A. McKenna, M. Hanna, E. Banks, A. Sivachenko, K. Cibulskis, A. Kernytsky, K. Garimella, D. Altshuler, S. Gabriel, M. Daly, M. A. DePristo, The Genome Analysis Toolkit: a MapReduce framework for analyzing next-generation DNA sequencing data. Genome Res. 20, 1297–1303 (2010).

78. H. Li, B. Handsaker, A. Wysoker, T. Fennell, J. Ruan, N. Homer, G. Marth, G. Abecasis, R. Durbin, 1000 Genome Project Data Processing Subgroup, The Sequence Alignment/Map format and SAMtools. Bioinformatics. 25, 2078–2079 (2009).

79. B. L. Browning, Y. Zhou, S. R. Browning, A One-Penny Imputed Genome from Next-Generation Reference Panels. Am. J. Hum. Genet. 103, 338–348 (2018).

80. 1000 Genomes Project Consortium, A. Auton, L. D. Brooks, R. M. Durbin, E. P. Garrison, H. M. Kang, J. O. Korbel, J. L. Marchini, S. McCarthy, G. A. McVean, G. R. Abecasis, A global reference for human genetic variation. Nature. 526, 68–74 (2015).

81. G. Barnes, State Formation in Korea: Emerging Elites (Routledge, 2013).

82. K. Taesik, The Cultural Characteristics of Korea’s Ancient Kaya Kingdom. International Journal of Korean History. 8, 169–221 (2005).

83. K. M. Wells, Korea: Outline of a Civilisation (BRILL, 2015).

84. D. Baker, Y. Ch’oe, H. H. W. Kang, H.-K. Kim, P. H. Lee, Sourcebook of Korean Civilization: From Early Times to the Sixteenth Century (Volume I) (Columbia University Press, 1993).

85. J. Ryan, G. Barnes, in *Encyclopaedia of the History of Science, Technology, and Medicine in Non-Western Cultures*, H. Selin, Ed. (Springer Netherlands, Dordrecht, 2014), pp. 1–16.

86. Kyungsung University Museum, The Daeseong-dong Cemetery in Gimhae I, II and III (2000).

87. I. U. Kang, Dictionary of Korean Archaeology 2014. Dictionary of Korean Archaeology (2014) (available at https://www.academia.edu/40681250/Dictionary_of_Korean_Archaeology_2014_Buyeo%E6%89%B6%E4%BD%99_Okcho%E6%B2%83%E6%B2%AE_Sushen%E8%82%85%E6%84%BC_Yilou%E9%82%91%E5%A9%81).

88. Gimhae National Museum of Korea, Yuha-ri site (Gimhae National Museum of Korea, Korea, 2012).

89. U. Buikstra, JE Buikstra, DH Ubelaker. Standards for data collection from human skeletal remains (1994).

90. S. Black, L. Scheuer, Age changes in the clavicle: From the early neonatal period to skeletal maturity. Int. J. Osteoarchaeol. 6, 425–434 (1996).

91. bamUtil (Github; https://github.com/statgen/bamUtil).

92. H. Li, A statistical framework for SNP calling, mutation discovery, association mapping and population genetical parameter estimation from sequencing data. Bioinformatics. 27, 2987–2993 (2011).

93. P. Danecek, A. Auton, G. Abecasis, C. A. Albers, E. Banks, M. A. DePristo, R. E. Handsaker, G. Lunter, G. T. Marth, S. T. Sherry, G. McVean, R. Durbin, 1000 Genomes Project Analysis Group, The variant call format and VCFtools. Bioinformatics. 27, 2156–2158 (2011).

94. S. R. Browning, B. L. Browning, Rapid and accurate haplotype phasing and missing-data inference for whole-genome association studies by use of localized haplotype clustering. Am. J. Hum. Genet. 81, 1084–1097 (2007).

